# DYRK2 activates heat shock factor 1 promoting resistance to proteotoxic stress in triplenegative breast cancer

**DOI:** 10.1101/633560

**Authors:** Rita Moreno, Sourav Banerjee, Angus W. Jackson, Jean Quinn, Gregg Baillie, Jack E. Dixon, Albena T. Dinkova-Kostova, Joanne Edwards, Laureano de la Vega

**Affiliations:** Division of Cellular Medicine, School of Medicine, University of Dundee, Scotland, UK; Department of Pharmacology, University of California San Diego, La Jolla, CA 92093-0721; Unit of Gastrointestinal Oncology and Molecular Pathology, Institute of Cancer Sciences, College of Medical, Veterinary, and Life Sciences, University of Glasgow, Glasgow, United Kingdom; Department of Cellular and Molecular Medicine, University of California San Diego, La Jolla, CA 92093, USA; Department of Chemistry and Biochemistry, University of California San Diego, La Jolla, CA 92093, USA

**Keywords:** DYRK2, Proteotoxic stress, HSF1, Triple-Negative Breast Cancer, HSP70, non-oncogene addiction

## Abstract

To survive aneuploidy-induced proteotoxic stress, cancer cells activate the proteotoxic-stress response pathway, which is controlled by heat shock factor 1 (HSF1). This pathway supports cancer initiation, cancer progression and chemoresistance and thus is an attractive target. As developing HSF1 inhibitors is challenging, the identification and targeting of upstream regulators of HSF1 presents a tractable alternative strategy. Here we demonstrate that in triple negative breast cancer (TNBC) cells, the dual-specificity tyrosine-regulated kinase 2 (DYRK2) phosphorylates HSF1, promoting its nuclear stability and transcriptional activity. Thus, DYRK2 depletion reduces HSF1 activity and sensitises TNBC cells to proteotoxic stress. Importantly, in tumours from TNBC patients, DYRK2 levels positively correlate with active HSF1 and associates with poor prognosis, suggesting that DYRK2 could be promoting TNBC. In agreement with this, DYRK2 depletion reduces tumour growth in a TNBC xenograft model. These findings identify DYRK2 as both, a key modulator of the HSF1 transcriptional program, and a potential therapeutic target.

## Introduction

Approximately 90% of solid human tumours and 75% of hematopoietic cancers exhibit some degree of aneuploidy^1^. In the majority of cases aneuploidy results from chromosomal instability (CIN), which produces the high level of genetic diversity that tumours require in order to evolve and develop. Evidence from yeast and mammalian systems has shown that aneuploid cells are generally less fit than their euploid counterparts^2, 3^. This is believed to be due to gene dosage imbalances, which cause an accumulation of excess and often misfolded proteins that must then be chaperoned and/or degraded in order to prevent proteotoxic stress^4, 5^. Consequently, proteotoxic stress represents a common problem that chromosomally unstable cells must overcome if they are to generate viable aneuploid progeny that are capable of malignant progression. In principle, this “aneuploidy tolerance” can be achieved by increasing either protein folding capacity or protein degradation, and therefore one might predict that positive regulators of these two processes are important for aneuploid cells to survive. This could explain why certain cancer cells are addicted to heat shock factor 1 (HSF1), the master regulator of proteotoxic stress responses, and to proteasome function. The identification of new ways to target these pathways, ideally simultaneously, might therefore provide new and generalised therapeutic opportunities to target cancer cells independently of their individual genetic lesions responsible for cancer initiation.

It is well established that HSF1 plays a critical role in many of the hallmarks of cancer cells and is important in tumour initiation^6^ and in promoting and maintaining cancer cell proliferation^6, 7^. High levels of HSF1 expression have been reported in a number of cancers^8^ and high levels of nuclear HSF1 are associated with poor outcome in breast cancer, hepatocellular carcinoma, and in endometrial carcinoma^9–11^ among others. Under basal conditions, HSF1 is maintained largely in the cytoplasm and upon proteotoxic stress, HSF1 is activated and translocates to the nucleus^12^. Once in the nucleus, HSF1 activates a comprehensive transcriptional program including a number of genes encoding heat shock proteins (HSPs), which function as molecular chaperones and act to protect cells against proteotoxic stress and apoptosis^13^. Of the HSPs, *HSP70*, is one of the best-characterised HSF1 targets, and is expressed at high levels in a variety of cancers, where its presence is correlated with growth, survival, invasion and resistance to therapy^14^. HSF1 activity and stability are tightly controlled by multiple post-translational modifications^15^. Among these, phosphorylation of serine 320 and serine 326 are associated with enhanced transcriptional activity, stability and nuclear accumulation of HSF1^16–18^. Although the sequence of modification events leading to HSF1 activation or the interplay among the different phosphorylation events are still unclear, the predominant view is that phosphorylation of S326 is both critical and dominant over any inhibitory phosphorylation events and is thus considered to be a hallmark of HSF1 activation^8^.

Recently, the dual-specificity tyrosine-phosphorylation-regulated kinase 2 (DYRK2) was shown to positively regulate proteasome activity^19^. Here we report that the CMGC kinase dual-specificity tyrosine-phosphorylation-regulated kinase 2 (DYRK2) phosphorylates and activates HSF1, thus increasing resistance to proteotoxic stress in triple negative breast cancer (TNBC) cells, and promoting tumour growth in TNBC xenograft models. We further report that the DYRK2-HSF1 link is relevant in clinical TNBC samples where DYRK2 protein levels correlate with active HSF1, and are associated with high rates of tumour recurrence and poorer patient survival. Taken together our data reveal that DYRK2 is a hitherto unknown and major regulator of the HSF1 pathway in vivo, which acts to promote TNBC growth and survival. TNBC is a tumour type characterised by genomic instability and aneuploidy, high rate of recurrence and metastases, and for which current therapeutic options are limited^20^. Small molecule inhibitors of DYRK2 may thus represent a novel approach to downregulate HSF1 activity and to increase the therapeutic options in treating TNBC.

## Results

### 1-DYRK2 phosphorylates HSF1

To test whether DYRK2 phosphorylates HSF1, we overexpressed DYRK2 and, by use of phosphospecific antibodies, we observed that the levels of endogenous HSF1 phosphorylated at S326 and S320 were increased (Fig. 1A); for this effect, the kinase activity of DYRK2 was required, as a kinase-dead version of DYRK2 (DYRK2-KD) did not induce HSF1 phosphorylation. To further test the role of DYRK2 kinase activity on HSF1 phosphorylation, we mutated the gatekeeper residue within DYRK2, creating an analogue-sensitive DYRK2 form (DYRK2-AS) that is selectively sensitive to PP1 inhibitors^21^. Using the DYRK2-AS form together with three different PP1 inhibitors, we observed that specific inhibition of DYRK2 activity, dose-dependently reduced the DYRK2-mediated phosphorylation of endogenous HSF1 (Fig. 1B). Additionally, harmine, a well characterised pan-specific DYRK inhibitor^22^, was also able to reduce the phosphorylation of endogenous HSF1 mediated by DYRK2 overexpression (Fig. S1A).

**Figure 1.**
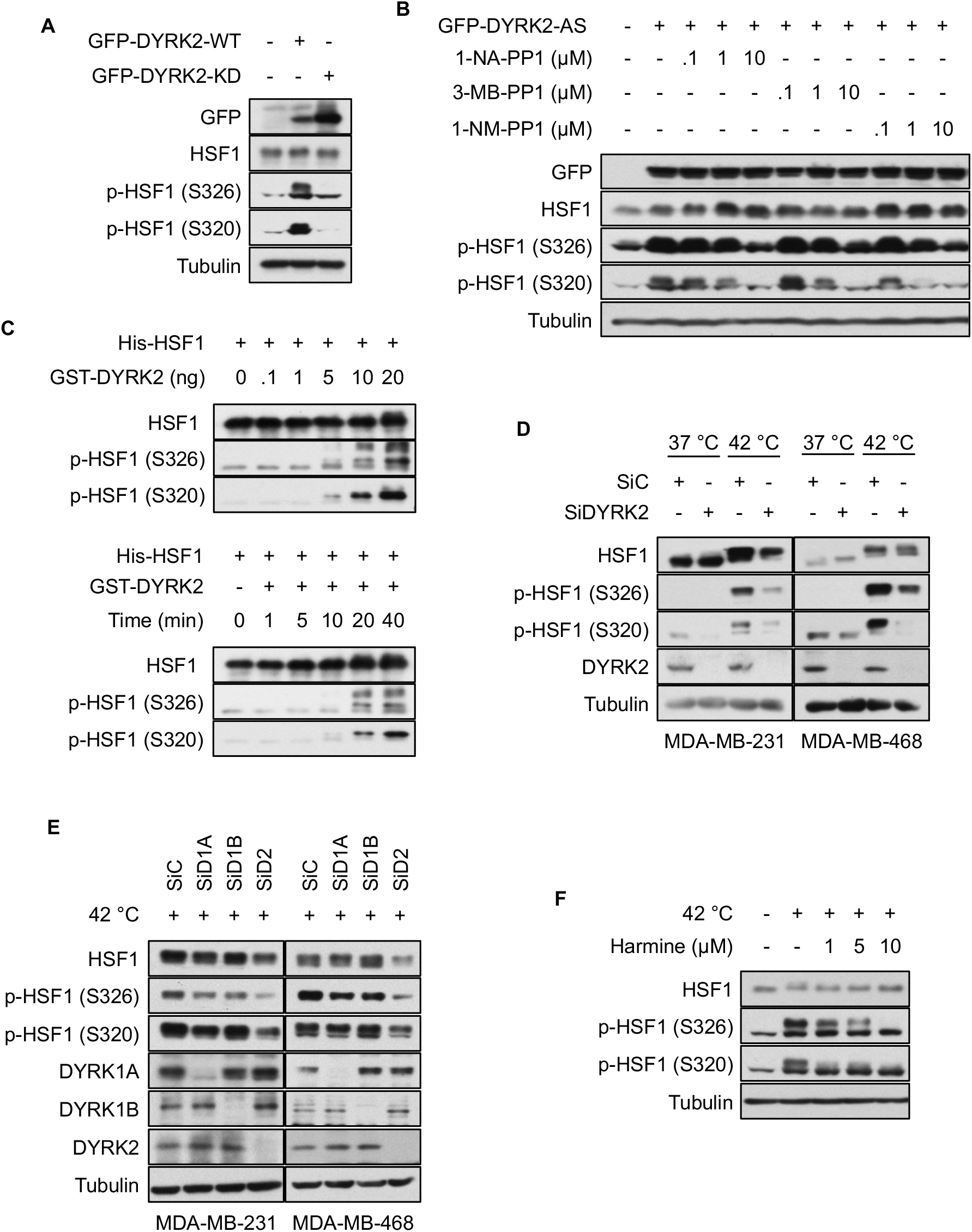
DYRK2 phosphorylates HSF1. **A)** 293T cells were transiently transfected to express GFP-tagged DYRK2 wild-type (WT) or a kinase dead (KD) version. After 48 hours, cells were lysed and the levels of endogenous HSF1 and phospho-HSF1 were analysed as indicated. **B)** 293T cells were transiently transfected with a GFP-tagged DYRK2 analog sensitive (AS) version. After 48 hours, cells were treated for a further 3 hours with increasing concentrations of three different PP1 inhibitors as indicated. Cells were lysed and the levels of endogenous HSF1 and phospho-HSF1 were analysed by western blot. **C)** *Upper panel,* purified recombinant His-HSF1 (1 μg) was incubated in kinase buffer with increasing concentrations of recombinant GST-DYRK2 at 30°C for 30 min. *Lower panel,* purified recombinant His-HSF1 (1 μg) was incubated with GST-DYRK2 (20 ng) at 30°C for various times as indicated. The reactions were terminated by the addition of SDS gel loading buffer, the proteins were resolved by SDS-PAGE, and the levels of phosphorylated HSF1 were analysed. **D)** MDA-MB-231 or MDA-MB-468 cells were transfected with either siControl or siDYRK2. After 48 hours, cells were either incubated at 42 °C for one hour or kept at 37 °C. Cells were lysed in SDS buffer and the levels of the indicated proteins were analysed by western blotting. **E)** MDA-MB-231 or MDA-MB-468 cells were transfected with either siControl, siDYRK1A, siDYRK1B or siDYRK2. 48 hours later cells were incubated at 42 °C for one hour. Cells were lysed in SDS buffer and the levels of the indicated proteins were analysed by western blotting. **F)** MDA-MB-231 cells were treated with increasing concentrations of harmine. One hour later, cells were incubated at 42 °C. After one hour, cells were lysed in SDS buffer and the levels of the indicated proteins were analysed by western blotting. See also Figure S1.

To address if HSF1 is a direct substrate for DYRK2, we performed an *in vitro* kinase assay using recombinant His-HSF1 and GST-DYRK2. In close agreement with the cell culture experiments, we found that DYRK2 phosphorylates His-HSF1 at S326 and S320 *in vitro* (Fig. 1C). Furthermore, incubation of recombinant GST-DYRK2-AS with a PP1 inhibitor inhibited the phosphorylation of His-HSF1 on both S326 and S320 (Fig. S1B). Mass spectrometry analysis of a sample taken at an early time point (5 minutes) from the *in vitro* reaction showed that HSF1 was further phosphorylated at S307, T323 and S363 (Fig. S1C).

Once we established that DYRK2 can phosphorylate HSF1 in cells and *in vitro,* we tested whether DYRK2 plays a role in HSF1 phosphorylation in response to proteotoxic stress in TNBC cell lines, using MDA-MB-231 and MDA-MB-468 as models. Using heat shock as a proteotoxic stress inducer, we showed that DYRK2 depletion leads to a reduction of HSF1 phosphorylation in response to heat shock (Fig. 1D). To test if this effect was specific for DYRK2, we silenced individually two other members of the DYRK family, DYRK1A and DYRK1B. We found that in our TNBC cell models, DYRK2 is the main DYRK member involved in HSF1 phosphorylation (Fig. 1E), although other members of the family can also modulate HSF1 to a lesser extent. This is also the case in other cancer cell lines (Fig. S1D). In agreement with the role of DYRKs in heat shock-mediated HSF1 phosphorylation, incubation with harmine dramatically impaired HSF1 phosphorylation in response to heat shock in TNBC cells (Fig. 1F). Although three other kinases (mTOR, MEK1 and p38) has been shown to be able to phosphorylate HSF1 at S326 ^18, 23, 24^, the strong effect observed with harmine, suggest that these kinases cannot compensate for the lack of DYRK activity in response to proteotoxic stress. To address their relevance in TNBC cells, we compared the effect of the DYRK inhibitor harmine, against the p38 inhibitor SB202190 and the mTOR inhibitor rapamycin at impairing HSF1 phosphorylation induced by heat shock (we did not include a MEK1 inhibitor as MEK1 does not regulate HSF1 in TNBC cell lines^24^). Our results demonstrate that of the inhibitors tested, the DYRK inhibitor is the most effective impairing HSF1 phosphorylation (Fig S1E).

Interestingly, in addition to reducing HSF1 phosphorylation, DYRK2 knockdown appeared to reduce the levels of HSF1 under heat shock conditions (Fig. 1D, 1E, and Fig. S1D), which could be a cause or a consequence of the lower HSF1 phosphorylation levels. However, acute (1 hour) harmine-mediated DYRK inhibition reduced HSF1 phosphorylation without affecting HSF1 levels (Fig. 1F) which, based on the HSF1 relatively long half-life (>6 hours)^18, 25^, suggests that reduction of HSF1 phosphorylation might precede or is uncoupled to HSF1 destabilisation.

### 2-DYRK2 interacts with HSF1 via two domains

Our results provide evidence to support a functional interaction between DYRK2 and HSF1. In order to test whether DYRK2 and HSF1 physically interact in cells, we performed co-immunoprecipitation. The interaction between the two proteins, which was not easily detectable at basal conditions, was increased upon exposure to proteotoxic stress (Fig. 2A and 2E). This weak interaction might reflect a transient binding between DYRK2 and HSF1, which is a common occurrence for kinases and their substrates. In order to map the interaction site(s), we used an *in vitro* interaction peptide array. By incubating recombinant GST-HSF1 with a membrane containing an array of overlapping peptides covering the entire DYRK2 protein, we identified two potential binding regions (Fig. 2B) within conserved aminoacid sequences (Fig. S2A). Binding region 1 (BR1) with sequence DDQGSYV, is located in a surface-exposed loop, and binding region 2 (BR2) with sequence TDA, falls within the unstructured C-terminus of DYRK2 (Fig. S2B). In order to validate the functional relevance of these two regions, we created DYRK2 constructs harbouring mutations either in BR1 or BR2, or in both simultaneously. Based on our results, BR1 appeared to be more important for the ability of DYRK2 to phosphorylate HSF1 (Fig. 2C). Furthermore, when both DYRK2 binding regions were mutated, HSF1 phosphorylation was strongly reduced in both S320 and S326, but notably, the ability of DYRK2 to phosphorylate SIAH2^26^, another known substrate, remained intact (Fig. 2D). This observation suggests that these two regions are not important for general DYRK2 kinase activity, but rather for its specific interaction with HSF1. As expected, the DYRK2 BR1/BR2 mutant (YV/AR-T/A) did not interact with HSF1 (Fig. 2E).

**Figure 2.**
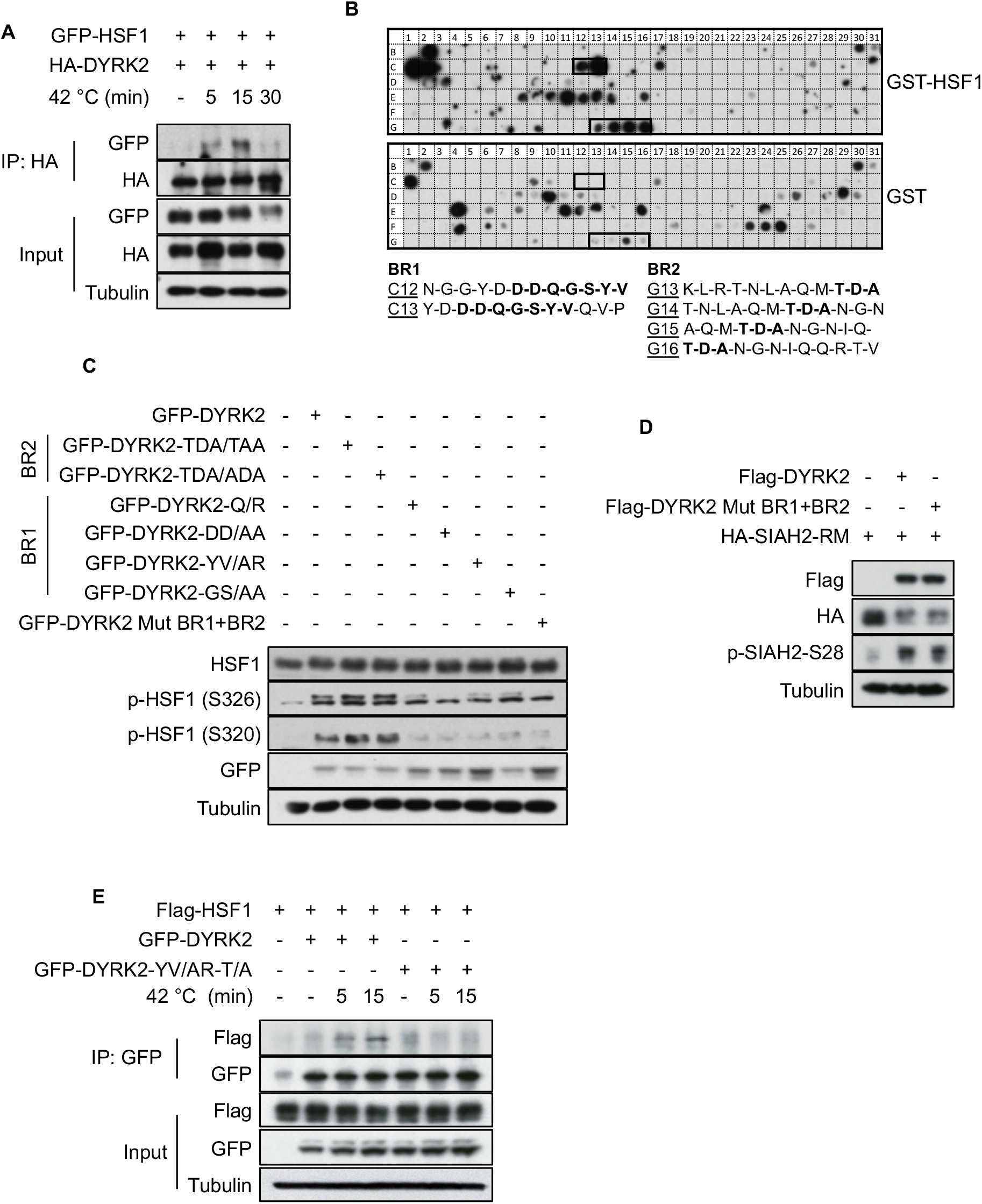
DYRK2 interacts with HSF1 via two domains. **A)** 293T cells were transfected with the indicated plasmids. After 48 hours, cells were incubated at 42 °C for the indicated periods of time. A fraction of the cell lysates was tested for the correct expression of the transfected proteins (Input), while the remaining extracts were used for immunoprecipitation with antiHA antibodies. After elution of bound proteins in 1 x SDS sample buffer, coprecipitated HSF1 was visualized by immunoblotting using an anti-GFP antibody as shown. **B)** A peptide array library covering the complete sequence of DYRK2 was incubated with recombinant GST-HSF1 protein *(upper panel)* or a GST control protein *(lower panel)* and the bound HSF1 protein was revealed by immunoblotting against GST as shown. The specific positive binding regions for HSF1 were indicated with a black box. Sequences of DYRK2 peptides within the binding region 1 (BR1) or binding region 2 (BR2) interacting with HSF1 are shown in bold. **C)** 293T cells were transiently transfected with the GFP-tagged versions of either DYRK2-WT or DYRK2 constructs harbouring mutations on the BR1 or BR2 as indicated, or a DYRK2 YV/AR-T/A mutant harbouring mutations in both regions (DYRK2 Mut BR1+BR2). After 48 hours, cells were lysed and the levels of endogenous HSF1 and phospho-HSF1 were analysed as indicated. **D)** 293T cells were transiently transfected with the Flag-tagged versions of either DYRK2-WT or DYRK2 Mut BR1+BR2, a DYRK2 mutant with mutations in both regions (DYRK2 YV/AR-T/A) together with an inactive form of SIAH2 (HA-SIAH2-RM). After 48 hours, cells were lysed and the levels of SIAH2 and phospho-SIAH2 were analysed as indicated. **E)** 293T cells were transfected with the indicated plasmids. After 48 hours, cells were incubated at 42 °C for the indicated periods of time. A fraction of the cell lysates was tested for the correct expression of the transfected proteins (input), while the remaining extracts were used for immunoprecipitation with anti-GFP antibodies. After elution of bound proteins in 1 x SDS sample buffer, coprecipitated HSF1 was visualized by immunoblotting using an anti-Flag antibody as shown. See also Figure S2

### 3-DYRK2 promotes HSF1 nuclear stability

As nuclear HSF1 is the active form of HSF1, we asked whether DYRK2 was affecting HSF1 nuclear levels. We tested in TNBC cells the role of DYRK2 on the nuclear levels of HSF1 at basal and after heat shock conditions by employing stable CRISPR-mediated knockout as well as siRNA-mediated knockdown. Both DYRK2 depletion strategies reduced HSF1 nuclear levels in basal conditions and after heat shock, as well as the levels of nuclear phospho-HSF1 in TNBC cells (Fig. 3A and Fig. S3A, S3B). This was also the case for other cancer cell lines (Fig. S3C and S3D). We used two independent gRNAs to knockout DYRK2 in the different TNBC cell lines, obtaining in both cases the same reduction in levels of nuclear HSF1 as with siDYRK2, suggesting a specific effect. Nevertheless, to further confirm the on-target effect of DYRK2 on HSF1 nuclear levels, we reconstituted DYRK2-KO cells with DYRK2, showing that the levels of nuclear HSF1 and phospho-HSF1 in DYRK2-KO cells were rescued (Fig. 3B). Moreover, DYRK2 overexpression resulted in increased HSF1 nuclear levels and phosphorylation (Fig. S3E). Interestingly, heat-shock induced a rapid nuclear stabilisation of DYRK2 (Fig. 3A, S3A, S3B and S3C), which further suggest that DYRK2 nuclear stabilisation is one of the mechanisms involved in the cellular response to proteotoxic stress in TNBC

**Figure 3.**
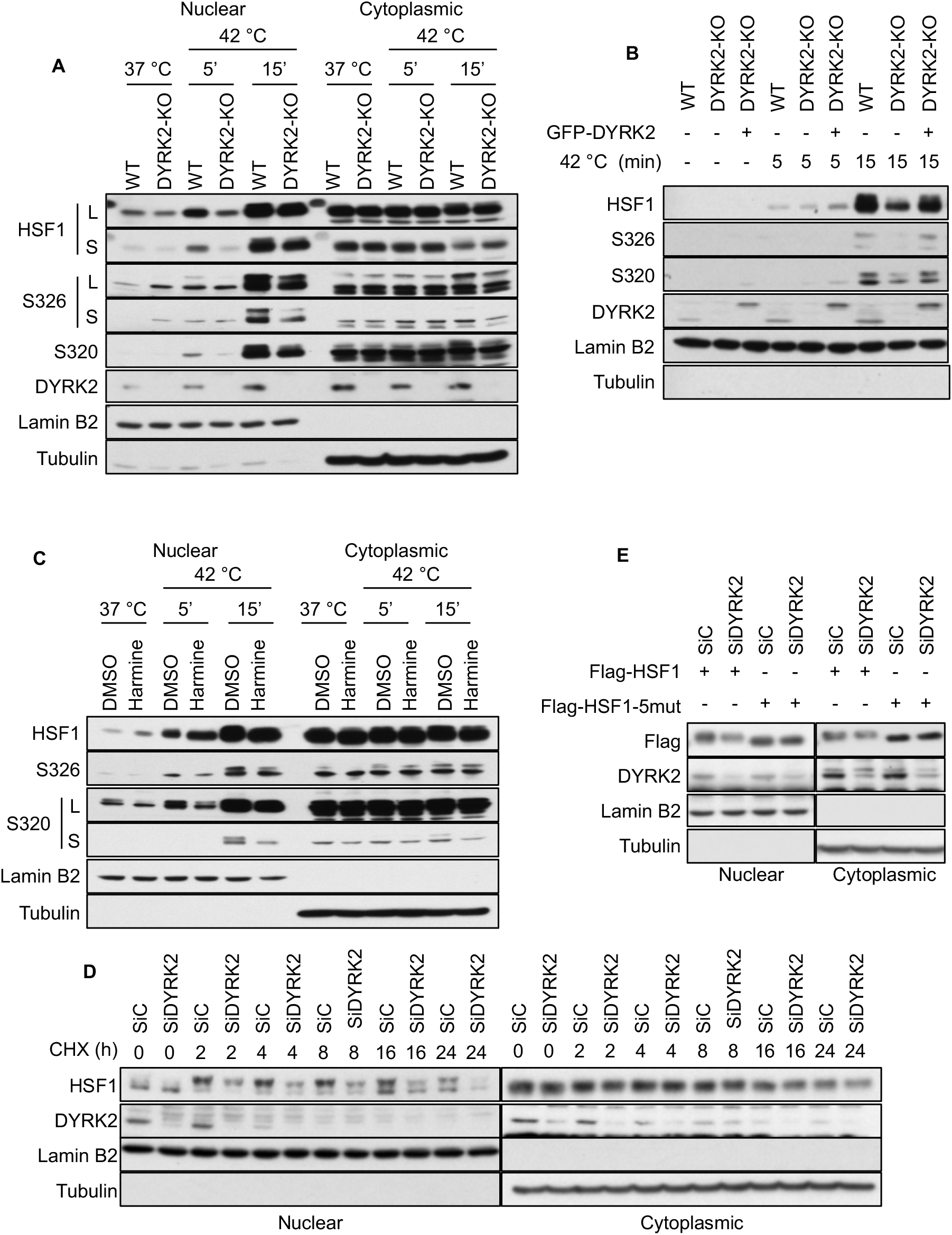
DYRK2 promotes HSF1 nuclear stability. **A)** Control (WT) and CRISPR-mediated DYRK2-KO MDA-MB-231 cells were incubated at 37 °C or at 42 °C for the indicated times. Nuclear and cytoplasmic fractions were analysed by western blot for the levels of the indicated proteins (L for long-exposure; S for short-exposure). **B)** DYRK2-KO 293T were transfected with either empty vector or with GFP-DYRK2. Control (WT), DYRK2-KO and DYRK2-GFP reconstituted DYRK2-KO cells were incubated at 37 °C or 42 °C for the indicated times. Cells were lysed, and nuclear fractions were analysed for the levels of the indicated proteins by western blotting. **C)** MDA-MB-231 cells were treated with either vehicle or 8 μM of Harmine. Two hours later, cells were incubated at 37 °C or 42 °C as indicated. After that, cells were lysed and nuclear and cytoplasmic fractions were analysed by western blotting (L for long-exposure; S for short-exposure). **D)** MDA-MB-231 cells were transfected with either siControl or siDYRK2, and 48 hours later cells were incubated with 20 μM of cycloheximide for the indicated times. Nuclear and cytosolic fractions were analysed for the levels of the indicated proteins by western blotting. **E)** 293T cells were transfected with either Flag-tagged HSF1 or with Flag-tagged HSF1 mutated in S307A, S320A, T323I, S326A, S363A (HSF1-5mut) in combination with siControl or siDYRK2 as indicated. After 48 hours, cells were exposed to 42 °C. After one hour, cells were lysed and nuclear and cytosolic fractions were analysed for the levels of the indicated proteins by western blotting. See also Figure S3

In agreement with our previous results, acute harmine-mediated DYRK inhibition led to reduction in nuclear HSF1 phosphorylation without affecting nuclear HSF1 levels (Fig. 3C). To assess the effect of DYRK2 on HSF1 stability, we compared the HSF1 nuclear and cytosolic levels upon cycloheximide treatment in control and DYRK2 knocked-down TNBC cells. In cycloheximide chase experiments, DYRK2 depletion reduced the levels of nuclear but not cytosolic HSF1, suggesting that DYRK2 controls HSF1 nuclear stability (Fig. 3D).

The positive effect of DYRK2 on HSF1 stability could be either dependent or independent of DYRK2-mediated HSF1 phosphorylation. We reasoned that as phosphorylation on S326 and on S320 are known to promote HSF1 stability and nuclear accumulation respectively^16, 18^, those two sites were most likely responsible for the positive effect of DYRK2 on HSF1 nuclear stability. Unexpectedly, our results showed that mutation of S326 and/or S320 were not sufficient to completely abolish the effect of DYRK2 on HSF1 stability (Fig. S3F). Nevertheless, additional mutation of the other three identified sites (S307, T323 and S363) significantly reduced the effect of DYRK2 on HSF1 stability (Fig. 3E). These results suggest the relevance of the combined phosphorylation on HSF1 for the effect of DYRK2 on HSF1 stability, although as some of those sites are also involved in HSF1 protein stability, we cannot completely rule out the participation of additional mechanisms unrelated to HSF1 phosphorylation.

### 4-DYRK2 affects the expression levels of the HSF1 target gene *HSP70*

Based on its positive effect on the levels of nuclear phosphorylated HSF1, we hypothesised that DYRK2 could be promoting HSF1 transcriptional activity. To test this hypothesis, we compared the levels of the prototypic HSF1 target gene, heat shock protein 70 *(HSP70),* in TNBC cells with and without DYRK2. DYRK2 knockout reduced significantly the induction of *HSP70* mRNA levels in response to heat shock (Fig. 4A and 4B) and in response to bortezomib, another inducer of proteotoxic stress (Fig. S4A). Similarly, HSF1 knockout strongly affected the levels of *HSP70,* whereas HSF1/DYRK2 double knockout did not reduce the *HSP70* mRNA levels any further, confirming that the effect of DYRK2 on the expression of *HSP70* is mediated by HSF1 (Fig. 4A). Importantly the reduction on *HSP70* levels observed in DYRK2-KO cells was recovered by reconstituting them with DYRK2 (Fig. 4C). In addition, harmine reduced the *HSP70* mRNA levels in a DYRK2-dependent manner in TNBC as well as in other cancer cell lines (Fig. S4B).

**Figure 4.**
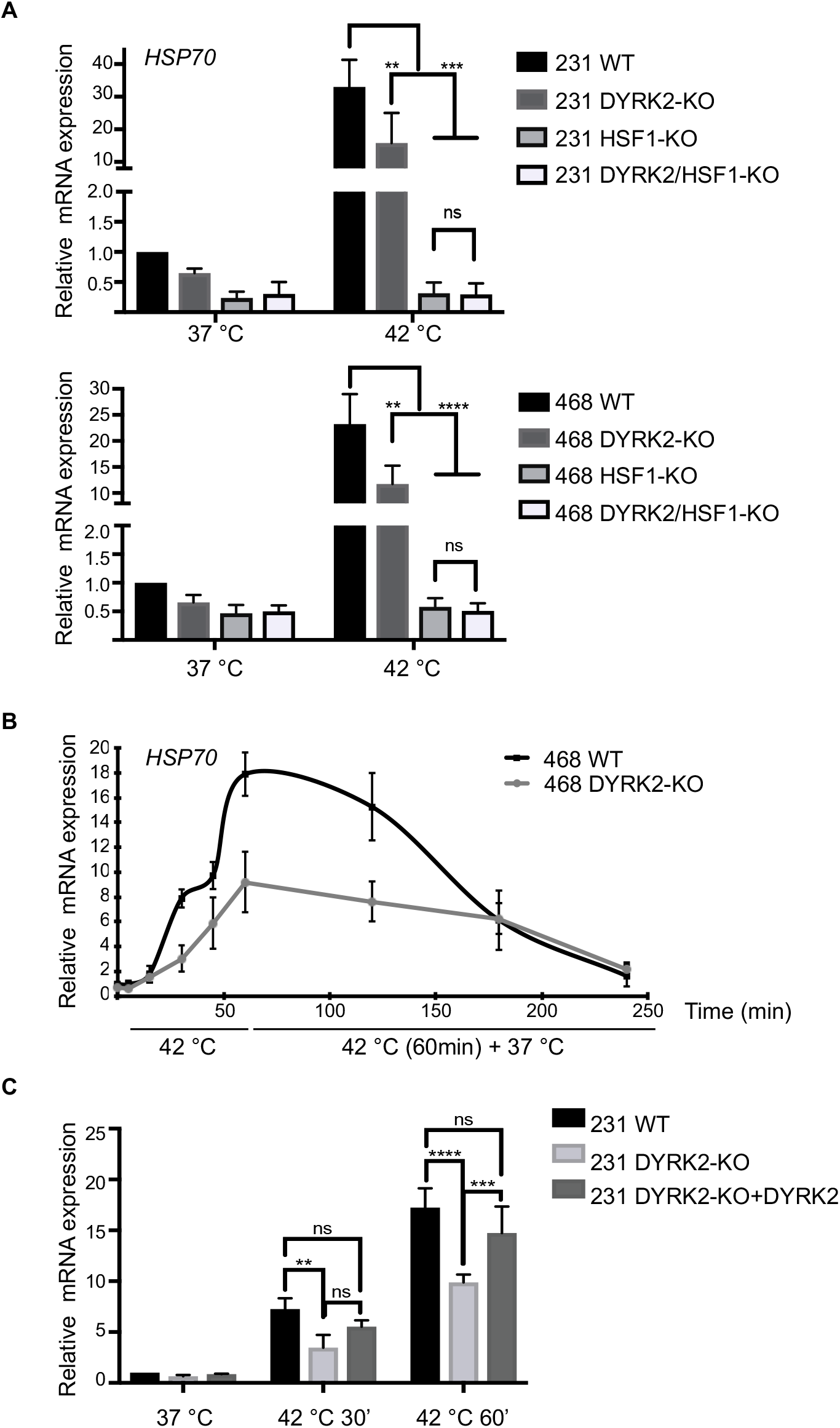
DYRK2 affects the expression levels of the HSF1 target gene *HSP70.* **A)** *Upper panel,* control (WT), DYRK2-KO, HSF1-KO or HSF1/DYRK2-KO MDA-MB-231 cells. *Lower panel,* control (WT), DYRK2-KO, HSF1-KO or HSF1/DYRK2-KO MDA-MB-468 cells. In both cases, cells were incubated at 37 °C or 42 °C for one hour. The mRNA levels for *HSP70 (HSPA1A)* were quantified using real-time PCR. The data were normalized using β-actin as an internal control. Data represent means ± SD (n=3) and are expressed relative to the control sample levels at 37 °C. *P ≤ 0.05, **P ≤ 0.01, ***P ≤ 0.001, ****P ≤ 0.0001. **B)** Control (WT) and DYRK2-KO MDA-MB-468 cells were incubated at 37 °C, at 42 °C for 5, 15, 30 or 60 minutes, or at 42 °C for 60 minutes plus recovery time at 37 °C (up to 180 minutes). After that, mRNA levels for *HSP70 (HSPA1A)* were quantified using real-time PCR. The data were normalized using β-actin as an internal control. Data represent means ± SD (n=3) and are expressed relative to the control samples at 37 °C. **C)** DYRK2-KO MDA-MB-231 were transfected with either empty vector or with GFP-DYRK2. Control (WT), DYRK2-KO and DYRK2-GFP reconstituted DYRK2-KO cells were incubated at 37 °C or 42 °C for the indicated times. After that, mRNA levels for HSP70 (HSPA1A) were quantified using real-time PCR. The data were normalized using β-actin as an internal control. Data represent means ± SD (n=3) and are expressed relative to the control samples at 37 °C. *P ≤ 0.05, **P ≤ 0.01, ***P ≤ 0.001, ****P ≤ 0.0001. See also Figure S4

### 5-DYRK2 reduces sensitivity to proteotoxic stress via HSF1

To assess the functional significance of the reduced HSF1 activity in DYRK2-impaired TNBC cells, we tested the effect of DYRK2 depletion on the sensitivity of these cells to proteotoxic stress. In agreement with our data on the importance of DYRK2 for HSF1 function and with the well-characterised protective role of HSF1 against proteotoxicity, DYRK2 depletion (both by siRNA and CRISPR-mediated knockout) sensitised TNBC cells to heat shock, as measured by levels of PARP cleavage, a marker of apoptosis (Fig 5A, 5B and Fig. S5A). The same result was observed in other cancer cell lines (Fig. S5B).

**Figure 5.**
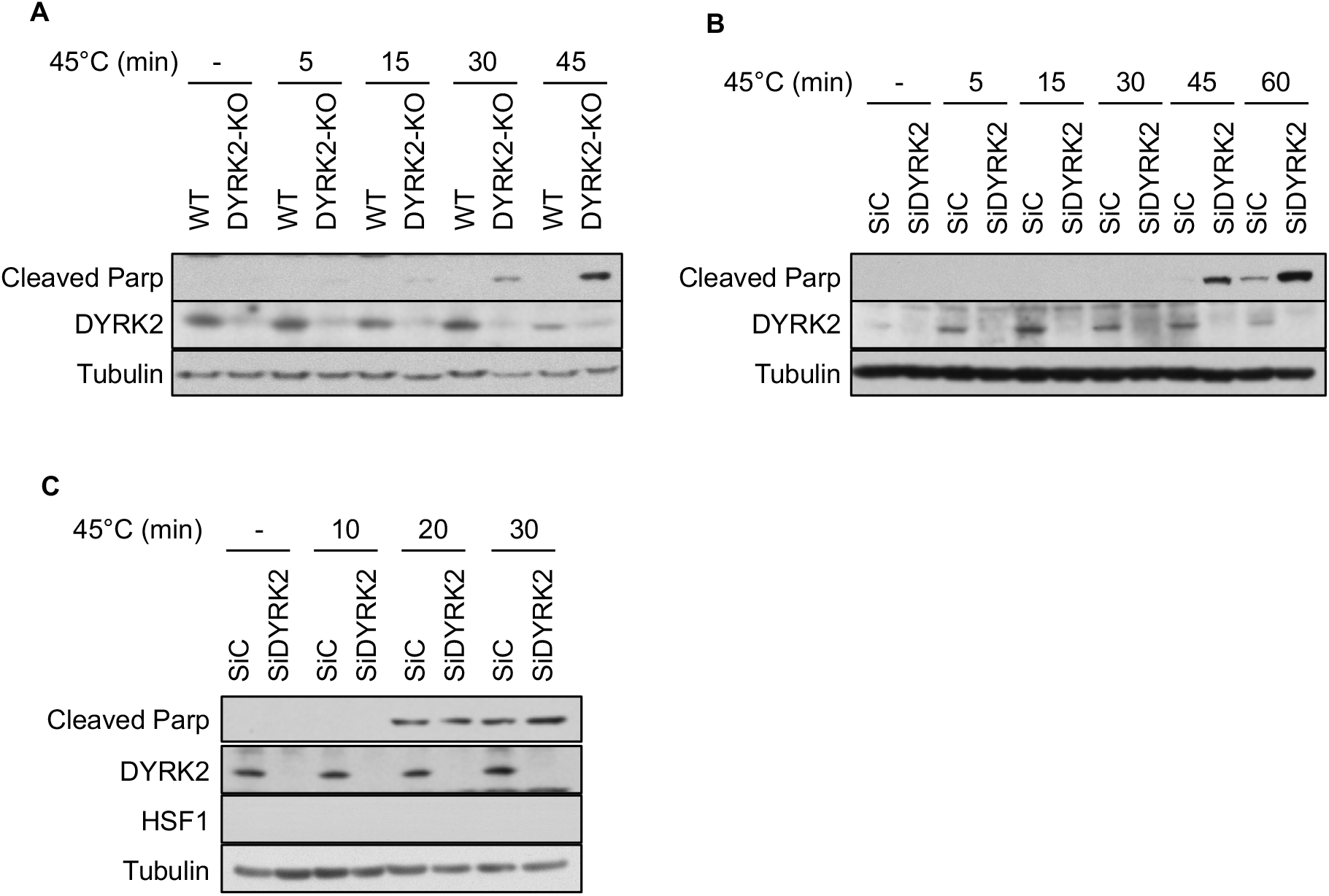
DYRK2 reduces sensitivity to proteotoxic stress via HSF1. **A)** Control (WT) and DYRK2-KO MDA-MB-231 cells were incubated at 37 °C (-) or at 45 °C for the indicated times, followed by recovery at 37 °C. On the next day, cells were lysed and the levels of apoptosis were analysed by western blotting using an antibody that recognises cleaved PARP. **B and C)** MDA-MB-231 (B) or HSF1-KO MDA-MB-231 (C) cells transfected with either siControl or siDYRK2 were incubated at 37 °C or at 45 °C for the indicated times, followed by recovery at 37 °C. On the next day, cells were lysed and the levels of apoptosis were analysed by western blotting using an antibody that recognises cleaved parp. See also Figure S5

As expected, HSF1-deficient cells were more sensitive to heat shock than wild-type cells. Importantly, the effect of DYRK2 depletion on the enhanced sensitivity to proteotoxic stress on wild-type cells was largely abolished in TNBC HSF1-KO cells (Fig. 5C), confirming the involvement of HSF1. In agreement, harmine sensitised TNBC cells to heat shock in a HSF1-dependent manner (Fig. S5C). Together, these results strongly suggest that DYRK2 has a role in the HSF1-mediated resistance to proteotoxic stress.

### 6-DYRK2 levels correlate with HSF1 nuclear levels, and associate with prognosis and tumour recurrence in tissue from TNBC patients

Next, to address if DYRK2 regulates HSF1 *in vivo,* we investigated whether in human TNBC tumours, the protein levels of DYRK2 correlate with levels of nuclear HSF1, which is a surrogate for active HSF1. To answer this question, we analysed by immunohistochemistry (IHC) the levels and localisation of both DYRK2 and HSF1 in 120 samples from TNBC patients (antibodies validation in figure S6A). When samples were divided into high and low expression groups for either HSF1 or DYRK2 (H-score: < or >145 for cytoplasmic and nuclear DYRK2; < or > 156 for cytoplasmic HSF1; and < or > 100 for nuclear HSF1), nuclear HSF1 positively correlated with nuclear DYRK2, but not cytoplasmic DYRK2 (Fig. 6A and S6B). Interestingly, when cytoplasmic HSF1 was analysed, no association was observed with DYRK2. Our data further support the conclusion that DYRK2 positively regulates HSF1 nuclear levels in tissue from TNBC patients.

**Figure 6.**
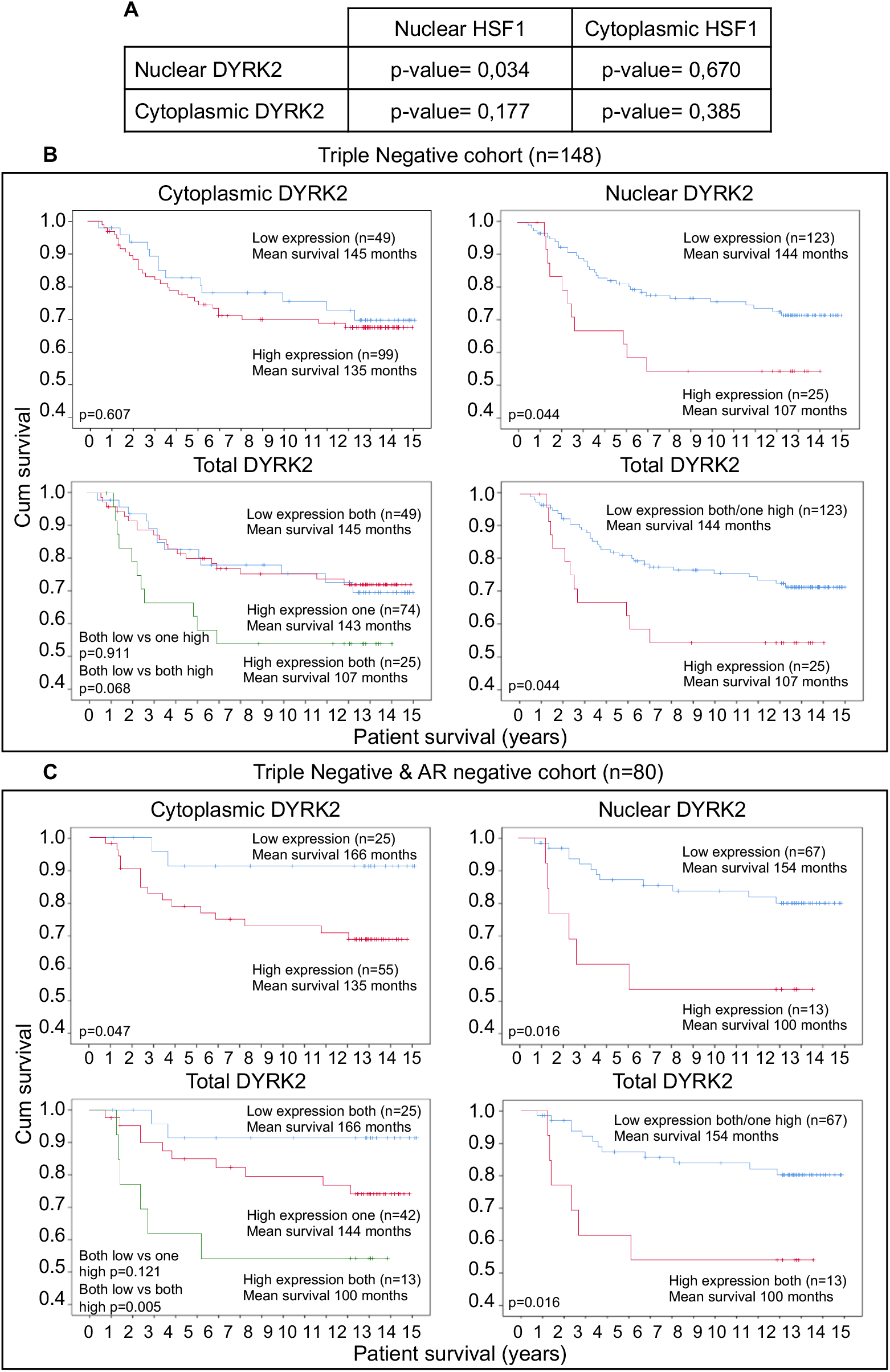
DYRK2 levels correlate with HSF1 nuclear levels and associates with poor prognosis and tumour recurrence in tissue from TNBC patients. **A)** Chi Square test correlations between nuclear/cytoplasmic HSF1, and nuclear/cytoplasmic DYRK2 in triple negative breast cancer. Significance p<0.05. **B)** Relationship between cytoplasmic, nuclear and total DYRK2 levels and cancer-specific survival in patients with triple negative invasive ductal breast cancer. “Low expression both” or “High expression both” means samples with low or high levels of both nuclear and cytoplasmic DYRK2; “High expression one” means samples with high levels of either nuclear or cytoplasmic DYRK2; “Low expression both/one high” means all the samples that do not have high levels of nuclear and cytoplasmic DYRK2. **C)** Relationship between cytoplasmic and nuclear DYRK2 levels and cancer-specific survival in patients with triple negative and AR negative invasive ductal breast cancer. See also Figure S6

Based on this novel correlation between DYRK2 and HSF1 in TNBC human samples and on the known association of nuclear HSF1 with poor prognosis, we hypothesised that high levels of nuclear DYRK2 would associate with poor prognosis in TNBC. To test this hypothesis, we utilised a cohort of 750 breast cancer samples that had available a previously constructed tissue microarray, including 148 TNBC (TMA information in Fig. S6C). Human tumour samples were stratified on the basis of high or low H-score (H-score: low<145 WHUs and high>145 WHUs). DYRK2 protein levels were assessed in 3 individuate TMA cores and mean WHU employed for analysis (representative figures with high and low DYRK2 levels in Fig. S6D). In the full cohort, DYRK2 levels did not associate with survival (Fig. S6E). However, in TNBC samples, high nuclear DYRK2 levels significantly associated with reduced cancer-specific survival (Fig. 6B) but not with overall survival (Fig. S6F). Notably, the strength of this association was highest in the TNBC subgroup of patients with ER, PR, HER2, and AR negative disease (Fig. 6C). In TN-AR negative samples, nuclear DYRK2 was observed as an independent factor in Cox Regression multivariate analysis when combined with other clinical parameters (Fig. S6G). Moreover, high DYRK2 levels were associated with shorter time to recurrence in both TNBC and TN-AR negative breast cancer (Fig. S6H).

These results establish DYRK2 as a potential prognostic factor and promising novel therapeutic target in TNBC, especially in the TN/AR-negative subgroup of patients, for whom there is no targeted therapy available. Interestingly, several of the commonly used TNBC cell models (including the two cell lines used in this study, MDA-MB-231 and MDA-MB-468) are also AR negative^27^, and thus there is full agreement between our cell culture and tissue analysis data.

### 7-The DYRK2-HSF1 axis promotes tumour growth in TNBC xenograft models

Based on our cell culture and tissue data, we wondered whether targeting this newly identified DYRK2-HSF1 link could affect tumour growth *in vivo.* To answer this question, and as there are no specific DYRK2 inhibitors available, we evaluated the tumour formation capacity of WT and HSF1-KO TNBC cells after DYRK2 knockdown. Tumours derived from TNBC DYRK2-deficient cells had significantly slower growth rates and smaller weight than those formed by DYRK2-proficient cells (Fig. 7A and 7B), confirming the role of DYRK2 in promoting TNBC tumour growth *in vivo.* Similarly, tumours derived from TNBC HSF1-KO cells grew more slowly and were lighter than those derived from WT cells. In this case, the effect of knocking down DYRK2 was less pronounced than in tumours from WT cells (Fig. 7a and 7B). Notably, the fact that tumours derived from TNBC HSF1-KO cells grew much more slowly than those from WT cells, made it difficult to unambiguously conclude that DYRK2 depletion does not affect tumour growth in HSF1-KO cells. Nonetheless, these experiments illustrate the importance of both DYRK2 and HSF1 for TNBC tumour growth and further show that, although HSF1 might not be the only mediator of the tumour-promoting activity of DYRK2, this kinase plays a major role in the growth of HSF1-proficient tumours. Overall, our data support the potential biological importance of the DYRK2-HSF1 axis in regulating cancer cell growth *in vivo.*

**Figure 7.**
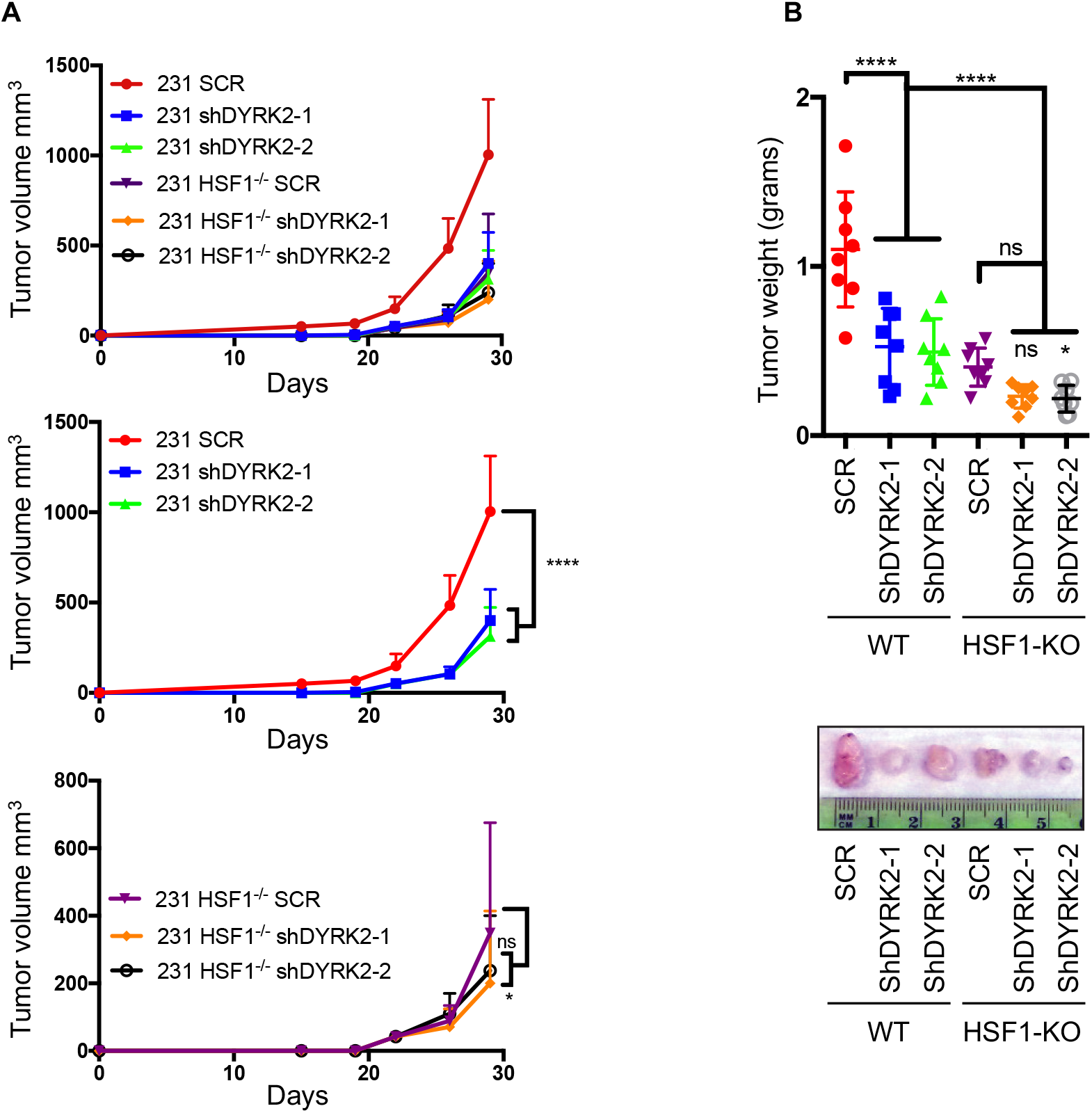
The DYRK2-HSF1 axis promotes tumour growth in TNBC xenograft models. **A)** DYRK2 was knocked down in control or HSF1-KO MDA-MB-231 cells using two independent shRNAs targeting DYRK2, along with a scrambled shRNA control. The cells were then injected subcutaneously into NSG^TM^ mice. Tumour volume was measured twice a week (n=8 per condition). **B)** Post 30 days of injection, tumours were resected, imaged and tumour weight was measured. ****p<0.0001, ***p<0.001, *p<0.05 (compared to scrambled control, twoway ANOVA, mean ± SD from n=8 mice each).

## Discussion

The HSF1 pathway represents an attractive therapeutic target as it allows cancer cells to adapt to and survive aneuploidy-induced proteotoxic stress, playing an important role in cancer initiation and also in cancer progression and chemoresistance^6, 8, 11, 28, 29^. However, developing specific HSF1 inhibitors has proven to be challenging, and thus, an alternative approach is to understand and to target HSF1 upstream regulatory pathways. In this study, we shed light on the regulation of the HSF1 pathway by the kinase DYRK2. We propose that DYRK2 is the major kinase controlling HSF1 activation in TNBC, and the first kinase described to phosphorylate HSF1 at both S326 and S320 sites. Remarkably, DYRK2 depletion significantly reduced HSF1 phosphorylation and its nuclear levels, and was sufficient to impair HSF1 response and promote TNBC apoptosis. This finding suggests that other kinases cannot compensate for the absence of DYRK2, and thus DYRK2 is critical for a full HSF1 activation in TNBC.

Interestingly, DYRK2 can phosphorylate HSF1 on at least five sites *in vitro,* and the mutagenesis studies show that only when all five sites are mutated, the effect of DYRK2 on HSF1 stability is lost, suggesting that those sites play a role *in vivo.* Although how these sites control HSF1 stability is not known, the fact that DYRK2 might phosphorylate sites associated with both positive (S326, S320) and negative (S307, S363) regulation of HSF1 might appear counter-intuitive. We propose that this conundrum could be explained by the current view that phosphorylation of S326 dominates over any other phosphorylation events. Thus, during heat shock, HSF1 undergoes both activating and inhibitory phosphorylation events, but as long as S326 is phosphorylated, the overall effect is HSF1 activation. Once S326 is dephosphorylated, the inhibitory phosphorylation events ensure that the HSF1 response is transient and does not persist.

We have shown that DYRK2 activates the pro-survival HSF1 pathway providing a support mechanism and a survival advantage to cancer cells. Importantly, this new link between DYRK2 and HSF1 appears to be relevant in TNBC patients as DYRK2 levels correlate positively with HSF1 nuclear levels, and negatively associates with cancer specific survival and time to recurrence, supporting a pro-tumoural role for DYRK2 in TNBC. Furthermore, we showed that DYRK2 predominantly affects nuclear HSF1, which is consistent with the observed increase in nuclear levels of DYRK2 in response to proteotoxic stress (Fig. 3A, 3B, S3A, S3B, S3C). This observation in cell lines, combined with the positive correlation between levels of nuclear DYRK2, nuclear HSF1 and association with patient prognosis observed in tumour samples suggest that the nuclear pool of DYRK2 is responsible for its HSF1-dependent tumour promoter role in TNBC.

Our results suggest that DYRK2 may be both a prognostic biomarker and a therapeutic target, which will be especially relevant in TN-AR negative breast cancer patients, for whom there is no targeted therapy available. The development of new specific DYRK2 inhibitors and their validation *in vivo* (i.e. using xenograft and genetic models) is needed to further substantiate this hypothesis. In this context, it is encouraging to see in our study the dramatic inhibitory effect of the DYRK inhibitor harmine on HSF1 activation and *HSP70* expression. Harmine has been shown to delay tumour growth in a genetic model of pancreatic carcinogenesis^30^, and although this drug inhibits various members of the DYRK family, our results support the relevance of the DYRK family as potential drug targets to impair cancer cell survival. In agreement, the drug curcumin inhibits DYRK2 activity and reduces TNBC tumour growth^31^, further highlighting the importance of developing new specific DYRK2 inhibitors which might have potential clinical relevance.

Our data, together with previous reports showing that DYRK2 controls proteasome activity^19^, presents DYRK2 as a major apical positive regulator of the two main pathways involved in alleviation of proteotoxic stress in cancer cells, and thus two main mechanisms by which aneuploid cells adapt, survive and become malignant. Thus, DYRK2 inhibitors by impairing both supporting mechanisms, might be an excellent tool to tackle not only TNBC cells, but also other aneuploid cancer cells that depend on such support mechanisms, independently on their genetic background.

## Acknowledgments

We thank Dr Stephen M. Keyse, Dr Kevin Hiom and Dr Adrian T. Saurin (University of Dundee) for critical reading and insightful comments on the manuscript. We would like to thank the Glasgow Biorepository for providing the tissue. We are extremely grateful to the Medical Research Institute of the University of Dundee, Cancer Research UK (C52419/A22869) (LV) and (CRUK/A18644) (ADK), Mary Kay Ash Breast Cancer grant 047.16 (JED) and UCSD Moores Cancer Centre Translational and Clinical Pilot Project award 2018 (JED) for financial support.

## Author Contributions

Conceptualization, R.M., A.D-K., and L.V.; Formal Analysis, J.E.; Investigation, R.M., S.B., A.J., GB and J.Q.; Resources, J.E.D. and J.E.; Writing – Original Draft, L.V.; Writing – Review & Editing, R.M., S.B., J.E., A.D-K. and L.V.; Visualization, R.M.; Funding Acquisition, J.E.D., J.E. and L.V.

## Declaration of Interests

The authors declare no competing interests.

## Material and Methods

### Cell culture

MDA-MB-231 were obtained from the culture collection at Public Health England. 293T, HeLa, and MDA-MB-468 cell lines were obtained from ATCC. All cell lines were grown in DMEM containing 10% FBS at 37°C and 5% CO2 and routinely tested for mycoplasma. Cells are passaged once 70-90% confluency is reached, and are maintained in culture for no more than 20 passages. Freshly thawed cells are passaged 2-3 times before used.

### Antibodies, plasmids and reagents

Anti-Flag Sigma-Aldrich; anti-HA (sc-805), anti-Lamin B2 (sc-56147) and anti-Tubulin (sc-8035), Santa Cruz Biotechnology; anti-GFP and (MRC Protein Phosphorylation and Ubiquitination Unit, University of Dundee, UK); anti-cleaved PARP (9546S) and anti-DYRK2 (8143), Cell Signalling; anti-HSF1 (ADI-SPA-901-D) Enzo Life Science; anti-phospo-HSF1-Ser 326 (ab115702), anti-phospo-HSF1-Ser 320 (ab76183) and anti GST-HRP (ab3416), Abcam. HRP-conjugated secondary antibodies (31430 and 31460) and Dynabeads™ Protein G (10004D), Life Technologies.

Antibodies recognizing Flag (F1804) were obtained from Sigma-Aldrich (Dorset, UK); anti-HA (sc-805), anti-Lamin B2 (sc-56147) and anti-Tubulin (sc-8035) were obtained from Santa Cruz Biotechnology (Dallas, Texas, USA). Anti-GFP anti-DYRK2 were obtained from the MRC Protein Phosphorylation and Ubiquitination Unit, (School of Life Science, University of Dundee, UK); anti-cleaved PARP (9546S) and anti-DYRK2 (8143) were obtained from Cell Signalling (MA, USA); anti-HSF1 (ADI-SPA-901-D) was obtained from Enzo Life Science (NY, USA); anti-phospo-HSF1-Ser 326 (ab115702), anti-phospo-HSF1-Ser 320 (ab76183) and anti GST-HRP (ab3416) were obtained from Abcam (Cambridge, UK). HRP-conjugated secondary antibodies (31430 and 31460) and Dynabeads™ Protein G (10004D) were obtained from Life Technologies (Carlsbad, California, USA).

Expression vectors for DYRK2-GFP, DYRK2-KD-GFP, DYRK2-Flag and SIAH2-RM-HA were a gift from Marco A. Calzado (University of Cordoba, Spain); HSF1-GFP have been already described^24^. All gRNAS were cloned into pLentiCRISPr-V2, which was a gift from Feng Zhang (Addgene plasmid # 52961). Point mutants were produced by conventional point mutagenesis. The DYRK2-analog sensitive was obtained by mutating the gatekeeper residue F228 to Alanine. His-HSF1 has already been described^24^ and GST-DYRK2 was obtained from the MRC Protein Phosphorylation and Ubiquitination Unit (School of Life Science, University of Dundee, UK). GFP-DYRK2 was cloned into the lentiviral 290-puro plasmid^32^ obtaining a puro resistant construct that was used for reconstitution experiments.

The siRNAs used against DYRK2 were the SMART pool: ON-Target Plus from Dharmacon (CO, USA). The shRNAs used to target DYRK2 (shDYRK2-1 gggtagaagcggtattaaa and shDYRK2-2 ggagaaaacgtcagtgaaa) were expressed from the pLL3.7 vector

Bortezomib, Santa Cruz Biotechnology; Harmine, Tocris Bioscience; PP1 inhibitors, Cayman Chemicals; SB202190 (1073), SYNkinase; Rapamycin (553210), Sigma-Aldrich Bortezomib (sc-217785) was obtained from Santa Cruz Biotechnology. Harmine (5075) was obtained from Tocris Bioscience (Bristol, UK), PP1 inhibitors (17860, 10954 and 13330) were obtained from Cayman Chemicals (MI, USA). the p38 inhibitor SB202190 (1073) was obtained from SYNkinase (3052, Australia) and the mTOR inhibitor Rapamycin (553210) was obtained from Sigma-Aldrich.

### Quantitative real time PCR (rt-qPCR)

RNA was extracted using RNeasy kit (QIAGEN). 500 ng of RNA per sample was reverse-transcribed to cDNA using Omniscript RT kit (QIAGEN) supplemented with RNase inhibitor (QIAGEN) according to the manufacturer’s instructions. Resulting cDNA was analysed using TaqMan Universal Master Mix II (Life technologies). Gene expression was determined using an Applied Biosystems 7300 Real-Time PCR system by the comparative ΔΔCT method. All experiments were performed at least in triplicates and data were normalized to the housekeeping gene β-actin. Probes were obtained from Applied Biosystems. When applicable, the differences between groups were determined by 2 way ANOVA. Analyses were performed using GraphPad Prism (GraphPad Software); a P value of <0.05 was considered significant. *P ≤ 0.05, **P ≤ 0.01, ***P ≤ 0.001.

### Cell transfections

On the day prior to transfection, cells were plated to the required cell density (70-90% confluency). Lipofectamine 2000 and Lipofectamine RNAiMAX (Invitrogen) were used for plasmid DNA and siRNA respectively. The plasmid DNA/siRNA and lipofectamine were individually diluted in Optimem (Gibco) and incubated for 10 minutes at room temperature. Diluted DNA/siRNA was added to the diluted Lipofectamine solution (1:1 ratio) and further incubated for 15 minutes. DNA-lipid complex was added to the cells and incubated overnight in a humidified incubator at 37°C and 5% CO2. The next morning, the medium was replaced with fresh medium and cells were incubated 36 hours more prior lysis.

### Lentivirus production

293T cells were transfected using Lipofectamine 2000 (Invitrogen) with the empty vector (pLL3.7) or the lentiviral shDYRK2-1 or shDYRK2-2 together with the packaging vectors (psPAX2 and pMD2.G) and cultivated in OptiMEM medium (Invitrogen). The next day the cells were further grown in DMEM complete medium and one day later the lentivirus-containing supernatant was collected, filtered and used to transduce cells.

### Cell lysis protocol and western blotting

Cells were washed and harvested in ice-cold phosphate-buffered saline (PBS) and lysed in either SDS buffer or RIPA buffer [50 mM Tris-HCl pH 7.5, 150 mM NaCl, 2 mM EDTA, 1% NP40, 0. 5% sodium deoxycholate, 0.5 mM Na3VO4, 50 mM NaF, 2 μg/ml leupeptine, 2 μg/ml aprotinin, 0.05mM pefabloc]. Cells directly lysed in SDS were boiled for two minutes, sonicated and boiled again for another 5 minutes. Cells lysed in RIPA buffer were sonicated and lysates were cleared by centrifugation for 15 minutes at 4°C. Protein concentration was established using the BCA assay (Pierce). Supernatant was mixed with SDS sample buffer and boiled for 5 minutes. Equal amounts of protein were separated by SDS-PAGE, followed by semidry blotting to a polyvinylidene difluoride membrane (Thermo Scientific). After blocking of the membrane with 5% (w/v) TBST nonfat dry milk, primary antibodies were added. Appropriate secondary antibodies coupled to horseradish peroxidase were detected by enhanced chemiluminescence using Clarity™ Western ECL Blotting Substrate (BIO-Rad).

### Subcellular fractionation (nuclear/cytoplasmic separation)

Cells were washed and harvested with ice-cold PBS. Pelleted cells were resuspended in 400μl of low-salt buffer A (10mM Hepes/KOH pH7.9, 10mM KCL, 0.1mM EDTA, 0.1mM EGTA, 1mM β-Mercaptoethanol). After incubation for 10 minutes on ice, 10μl of 10% NP-40 was added and cells were lysed by gently vortexing. The homogenate was centrifuged for 10 seconds at 13,200 rpm in a microfuge. The supernatant representing the cytoplasmic fraction was collected and the pellet containing the cell nuclei was washed 4 additional times in buffer A, then resuspended in 100μl high-salt buffer B (20mM Hepes/KOH pH7.9, 400mM NaCL, 1mM EDTA, 1mM EGTA, 1mM β-Mercaptoethanol). The lysates were sonicated and centrifuged at 4°C for 15 minutes at 13,200 rpm. The supernatant representing the nuclear fraction was collected. Protease and phosphatase inhibitors were freshly added to both buffers.

### Immunoprecipitation

Cells were washed with PBS and lysed in IP buffer (50 mM Hepes pH 7.5, 50 mM NaCl, 1% (v/v) Triton X-100, 2 mM EDTA, 10 mM sodium fluoride, 0.5 mM sodium orthovanadate, leupeptine (10 μg/ml), aprotinin (10 μg/ml), and 1 mM PMSF), followed by a sonication step. The extract was centrifuged and the supernatant was transferred to a new tube. The immunoprecipitation was performed with 2 μg of precipitating antibodies together with 50 μl of Dynabeads™ Protein G. Tubes were rotated for 30 min on a spinning wheel at 4°C. The immunoprecipitates were washed 3x with PBS/0,01% Tween-20 and eluted by boiling in 1 x SDS sample buffer. Equal amounts of protein were separated by SDS-PAGE.

### CRISPR-edited cell lines

The endogenous DYRK2 and/or HSF1 genes were knocked-out by transfecting cells with pLentiCRISPR-v2 (which codes for Cas9, and a puromycin cassette) containing gRNAs against the first exon of the short DYRK2 isoform or against the fourth exon of HSF1. For MDA-MB-231, HeLa and 293T DYRK2-KO cells the gRNA sequence used was GCTTGCCAGTGGTGCCAGAG and for MDA-MB-468 DYRK2-KO cells the target sequence was CGCTCACGGACAGATCCAGG. Additionally, we also tested some of our results in MDA-MB-231 cells in which we almost completely remove the DYRK2 ORF by using two gRNAs (N-term sequence GCTTGCCAGTGGTGCCAGAG and C-term sequence GAAGCTGAGCTAGAAGGTGG). For HSF1-KO cells the gRNA sequence used was AAGTACTTCAAGCACAACAA. Control cells were transfected with the empty pLentiCRISPRV2 vector. After transfection, cells were exposed to 2 μg/ml of puromycin for two days followed by a medium exchange. Surviving cells were clonally selected (in the case of control cells were used as pool population) by serial dilution, and positive clones were identified by genomic analysis and western blot. At least two clones for each cell line were used for the experiments.

### In vitro kinase assay

Bacterially expressed full length HSF1 His tagged was incubated with GST-DYRK2 in Kinase Buffer (25 mM TRIS pH 7.5, 5 mM β-glycerophosphate, 10 mM MgCl2 and freshly added 2mM DTT, 0.1 mM Na3VO4, 1 mM ATP) for different periods of time (5-60 min) at 30 °C. The reaction was inactivated by adding SDS loading buffer (50 mM Tris-HCl pH 6.8, 10%, SDS, 40% glycerin, 15% β-mercaptoethanol, 0.1 % bromophenol blue) and samples were boiled at 95 °C for 5 minutes and loaded in SDS-PAGE gels.

### In vitro peptide binding

Overlapping dodecapeptides covering the entire DYRK2 protein were spotted in an automated process on cellulose membranes by using Fmoc-protection chemistry (Proteomics facility, CNB-CSIC, Spain). The membrane was blocked overnight in non-fat milk in TBS buffer and incubated for 6 h with 60 nmol of recombinant GST-HSF1 or GST proteins as a control. After extensive washing, anti-GST antibody coupled to horseradish peroxidase was added for 1 hour. After washing, blots were revealed by enhanced chemiluminescence using Clarity™ Western ECL Blotting Substrate (BIO-Rad).

### Mass Spectrometry

Purified His-HSF1 (1ug) was incubated with GST-DYRK2 (20 ng) in 20 uL buffer containing 25mM Tris-HCl (pH7.5), 5 mM MgCl2, and 1 mM ATP for 5 minutes. The mixtures were reduced, alkylated and digested with trypsin (1:100) prepared in 100mM TEAB buffer (pH 8.5). The mixture was incubated at 37°C overnight before 8 uL of 10mM EDTA (in 0.1% TFA) was added. The peptide mixture was then subjected to nano-LC-MS/MS analysis according to a previous report^33^. Database search was carried out using Peaks 7.0 against UniProt-human (version 2018-05-28). False discovery rate was set at 1% (for peptide spectrum matches). The non-modified and modified (methionine oxidation and phosphorylation) peptides from HSF1 detected are shown in blue lines. The putative oxidation sites are labelled in yellow and phosphorylation sites in red.

### Xenograft assays

Female NSG^TM^ or Homozygous Foxn1nu (J:NU 007850) (Jackson Labs, USA) mice were housed and maintained at the University of California-San Diego (UCSD) in full compliance with policies of the Institutional Animal Core and Use Committee (IACUC). DYRK2 was knocked down in MDA-MB-231 (parental or HSF1 KO) cells with two independent shRNA oligoes as described previously^19^. MDA-MB-231 parental or HSF1 KO with either scrambled control or DYRK2 knock-down shRNA (1 or 2) were counted and suspended in sterile phosphate buffer saline containing 50% Matrigel (BD Bioscience). 300,000 cells (100 μl) were injected subcutaneously into the neck of each mouse. Tumour dimensions were measured twice per week using a digital caliper and tumour volume was calculated as (length × width^2^)/2. After the indicated days post-injection, mice were euthanized and tumours were excised and weighed.

Eight mice per group was the standard sample size for tumour xenograft experiments as defined by statistical power analyses (80-90% power; p<0.05) carried out using R packages. The investigators were not blinded to allocation during experiments and outcome assessment. Data were analysed using Graphpad Prism statistical package. All results are presented as mean ± SD unless otherwise mentioned

### Patient cohort and Tissue microarrays

Patients presenting with invasive ductal breast cancer at Glasgow Royal Infirmary, Western Infirmary and Stobhill Hospital between 1995 and 1998, with formalin-fixed paraffin embedded blocks of the primary tumour available for evaluation were studied (n =850). The study was approved by the Research Ethics Committee of the West Glasgow University Hospitals NHS Trust. Clinicopathological data including age, tumour size, tumour grade, lymph node status, type of surgery and use of adjuvant treatment (chemotherapy, hormonal based therapy and/ or radiotherapy) were retrieved from the routine reports. Tumour grade was assigned according to the Nottingham Grading System. ER and PR status were assessed on tissue microarrays (TMAs) using immunohistochemistry (IHC) with Dako (Glostrup, Denmark) ER antibody and Leica (Wetzlar, Germany) PR antibody. Her2 status were assessed as described^34^.

### Tissue analysis

IHC was conducted in triplicate on tissue microarrays (TMAs). Prior to this, antibody specificity was confirmed using proficient and deficient cell lines (either knockdown or knockout cell lines) (Fig S6A). TMA sections (2.5μm) were dewaxed by immersion in histoclear then rehydrated using a series of alcohols. Heat induced antigen retrieval was performed in a solution of either Tris-EDTA pH9 (DYRK2) or citrate buffer pH6 (HSF1) after which the sections were incubated in 3% hydrogen peroxide. Non-specific binding was blocked by incubation in 1.5% normal horse serum before being incubated with the optimum concentration of primary antibody overnight at 4°C. Antibody dilutions were prepared in antibody diluent (Agilent, London, UK). The primary antibodies and concentrations used are as follows; DYRK2 (Abgent AP7534a) at 1:200 and HSF1 (Cell Signaling 4256) at 1:500. Staining was visualized using ImmPRESS^TM^ and ImmPACT ^TM^DAB, (both Vector Labs). Tissue was counterstained using Harris Haematoxylin before being dehydrated and mounted using DPX. The stained TMA sections were scanned using a Hamamatsu NanoZoomer (Welwyn Garden City, Hertfordshire, UK) at x20 magnification. Visualization was carried out using Slidepath Digital Image Hub, version 4.0.1 (Slidepath, Leica Biosystems, Milton Keynes, UK). The weighted histoscore method was employed to assess protein expression [GB], with a second independent observer [JE] scoring 10% of the scores. Histoscores were calculated using the following formula: 0x % negative staining + 1x % of weakly positive staining + 2x % of moderately positive staining +3x % of strongly positive staining giving a range of 0 (minimum) to 300 (maximum). The results were considered discordant if the histoscores differed by more than 50 weighted histoscore units (WHU). Cytoplasmic and nuclear expression, if observed, was calculated separately.

## Supplementary Figure legends

**Figure S1.**
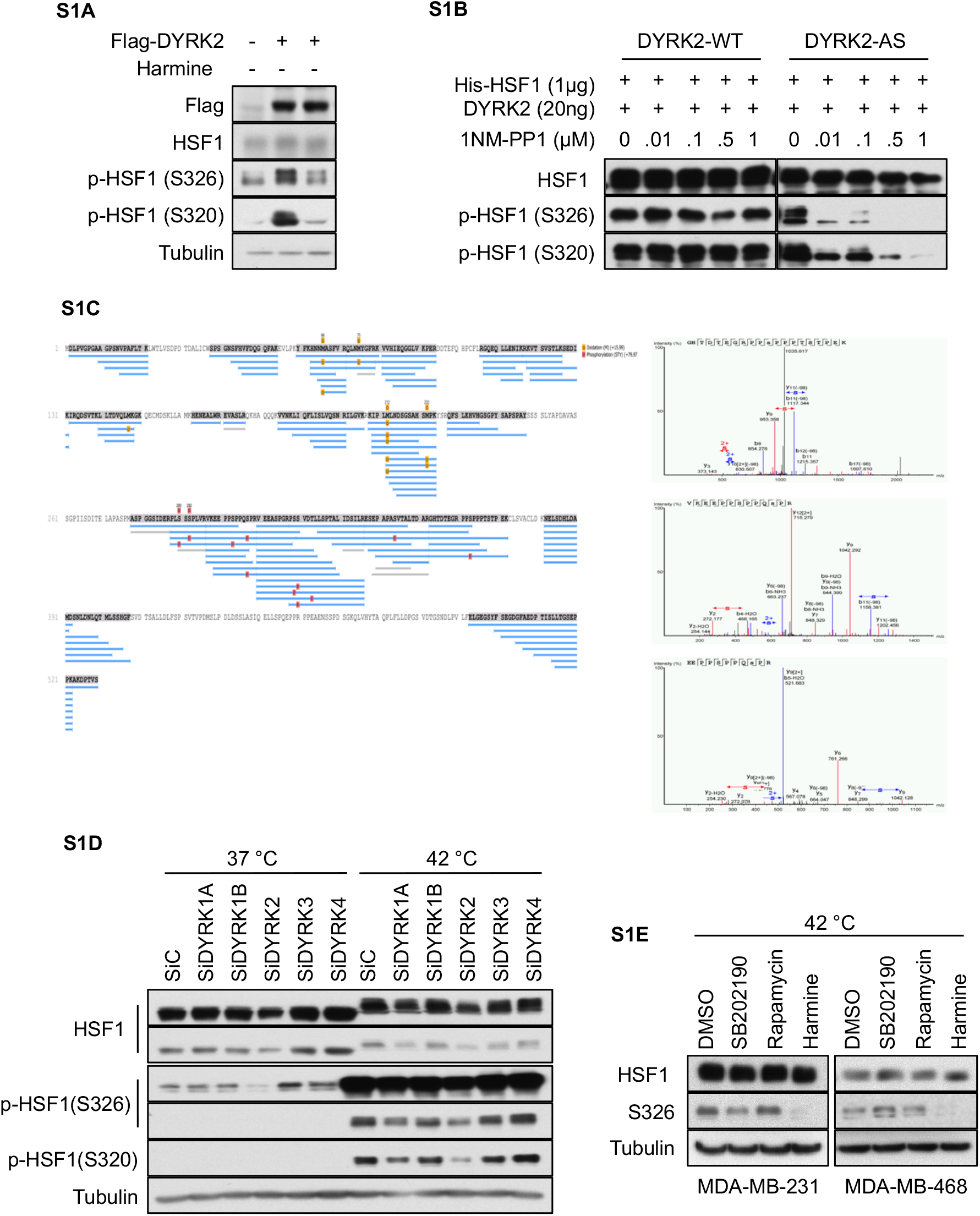
Related to Figure 1. DYRK2 phosphorylates HSF1. **A)** 293T cells were transiently transfected with empty vector or Flag-tagged DYRK2 as indicated. After 48 hours, cells were treated for a further 3 hours with vehicle or harmine (8 μM). Cells were lysed and the levels of endogenous HSF1 and phospho-HSF1 were analysed as indicated. **B)** 20 ng of recombinant GST-DYRK2-WT *(left panel)* or GST-DYRK2-AS *(right panel)* were incubated with His-HSF1 (1 μg) and increasing concentrations of 1NM-PP1 in kinase buffer without ATP. After 5 minutes, ATP was added and the reaction incubated at 30°C for 30 min. The reactions were terminated by the addition of SDS gel loading buffer, the proteins were resolved by SDS-PAGE, and the levels of phosphorylated HSF1 were analysed. **C)** Nano-LC-MS/MS analysis of His-HSF1 incubated with GST-DYRK2 for 5 minutes. Database search was carried out using Peaks 7.0 against UniProt-human(version 2018-05-28). False discovery rate was set at 1% (for peptide spectrum matches). The non-modified and modified (methionine oxidation and phosphorylation) peptides from HSF1 detected are shown in blue lines. The putative oxidation sites are labled in yellow and phosphorylation sites in red. **D)** HeLa cells were transfected with either siControl or siRNAs against all members of the DYRK family. 48 hours later cells were incubated at 37 °C or 42 °C for one hour. Cells were lysed in SDS buffer and the levels of the indicated proteins were analysed by western blotting. **E)** MDA-MB-231 and MDA-MB-468 cells were treated with either DMSO, the p38 inhibitor SB202190 (10 μM), the mTOR inhibitor rapamycin (30 nM) or harmine (10 μM). One hour later, cells were incubated at 42 °C. After one hour, cells were lysed in SDS buffer and the levels of the indicated proteins were analysed.

**Figure S2.**
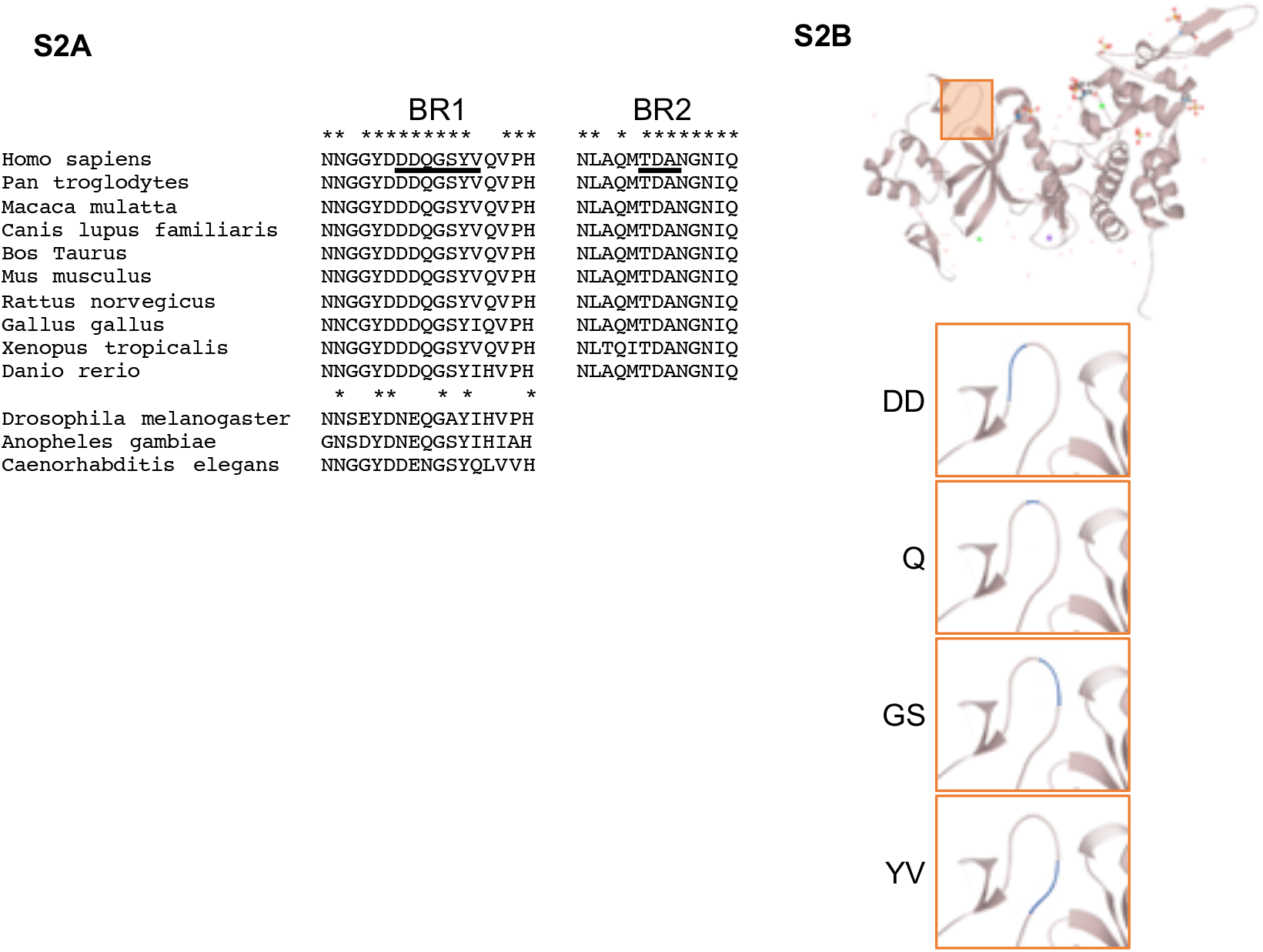
Related to Figure 2. DYRK2 interacts with HSF1 via two domains. **A)** Species alignment for the DYRK2 sequence containing the two identified binding regions, BR1 and BR2 (underlined). Asterisks mark conserved residues. **B)** Representation of the 3D structure of DYRK2 obtained for UniProt. The square focused on the DDQGSYV region and shows in blue the position of the different residues.

**Figure S3.**
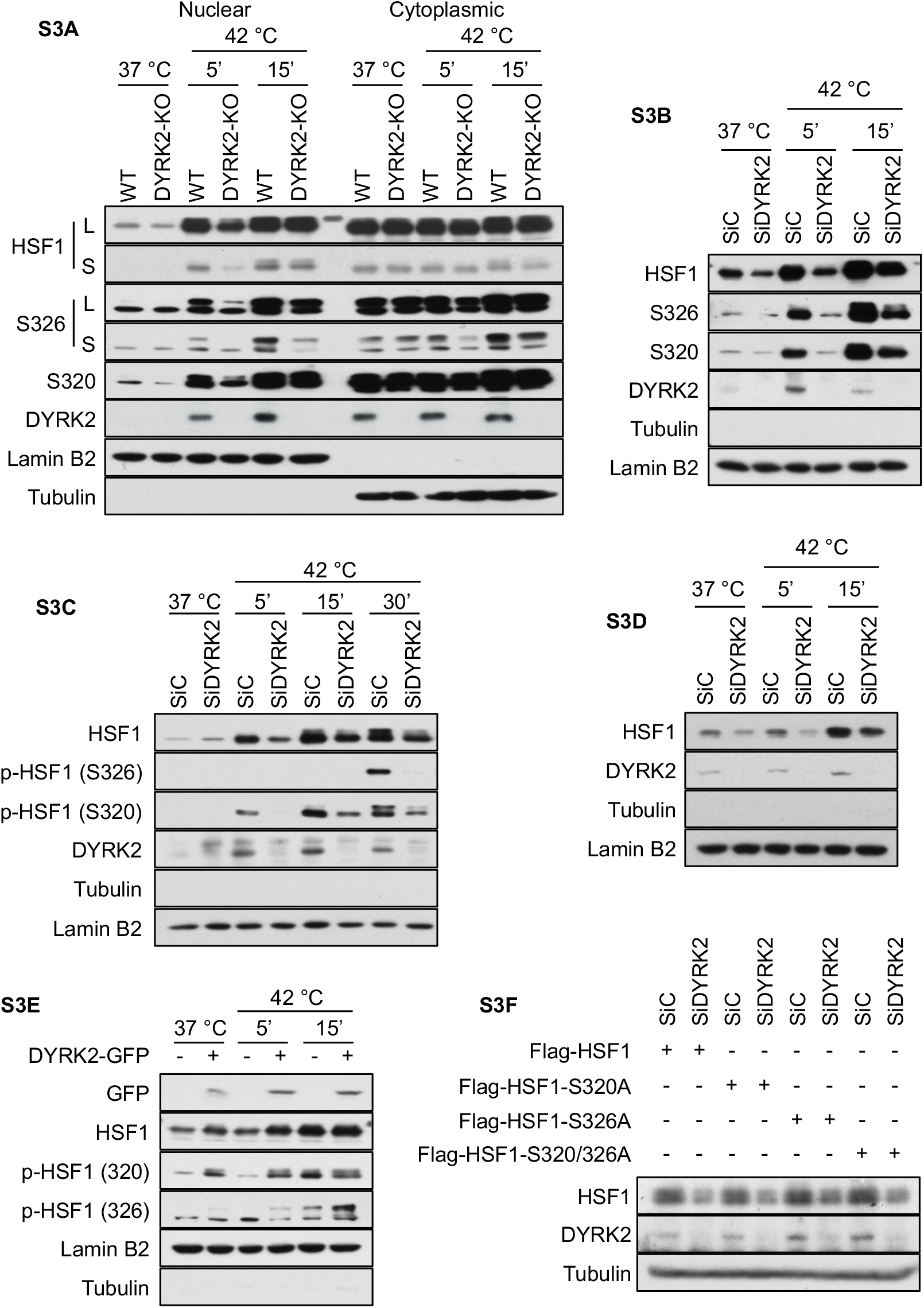
Related to Figure 3. DYRK2 promotes HSF1 nuclear stability. **A)** Control (WT) and CRISPR-mediated DYRK2-KO MDA-MB-468 cells were incubated at 37 °C or at 42 °C for the indicated times. Nuclear and cytosolic fractions were analysed by western blot for the levels of the indicated proteins (L for long exposure; S for short exposure). **B, C and D)** MDA-MB-468 cells (B), HeLa cells (C) or 293T cells (D) were transfected with either siControl or siDYRK2. 48 hours later cells were incubated at 37 °C or at 42 °C for the indicated times. Nuclear fractions were analysed by western blot for the levels of the indicated proteins. **E)** 293T cells were transfected with either empty vector or GFP-DYRK2. After 48 hours, cells were incubated at 37 °C or 42 °C for the indicated periods of time. Nuclear fractions were analysed by western blot for the levels of the indicated proteins. **F)** 293T cells were transfected with either Flag-tagged HSF1 or with the indicated Flag-tagged mutant HSF1 in combination with siControl or siDYRK2. After 48 hours, cells were exposed to 42 °C. After one hour, cells were lysed and analysed for the levels of the indicated proteins by western blotting

**Figure S4.**
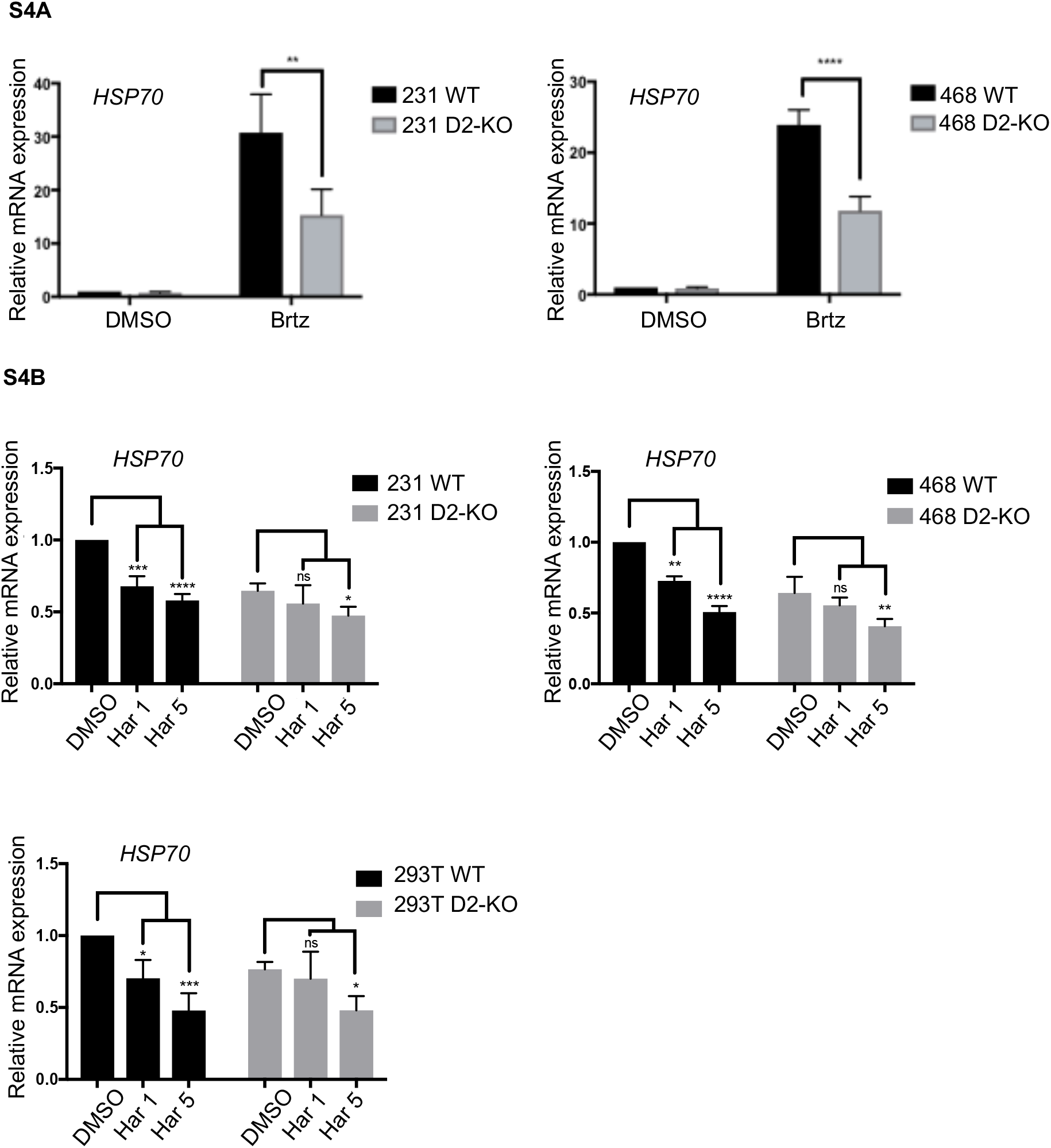
Related to Figure 4. DYRK2 affects the expression levels of the HSF1 target gene *HSP70.* **A)** Control (WT) and DYRK2-KO MDA-MB-231 cells *(left panel),* or Control (WT) and DYRK2-KO MDA-MB-468 cells *(right panel),* were incubated with either vehicle or 20nM Bortezomib (Brtz) for four hours. The mRNA levels for *HSP70 (HSPA1A)* were quantified using real-time PCR. The data were normalized using β-actin as an internal control. Data represent means ± SD (n=3) and are expressed relative to the control samples treated with vehicle. *P ≤ 0.05, **P ≤ 0.01, ***P ≤ 0.001, ****P≤ 0.0001. **B)** Control (WT) or DYRK2-KO MDA-MB-231 cells *(left panel),* or MDA-MB-468 cells *(right panel),* or 293T cells *(lower panel)* were incubated with increasing concentrations of harmine. After 2 hours, the mRNA levels for *HSP70 (HSPA1A)* were quantified using real-time PCR. The data were normalized using β-actin as an internal control. Data represent means ± SD (n=3) and are expressed relative to the control samples treated with vehicle. *P ≤ 0.05, **P ≤ 0.01, ***P ≤ 0.001, ****P≤ 0.0001.

**Figure S5.**
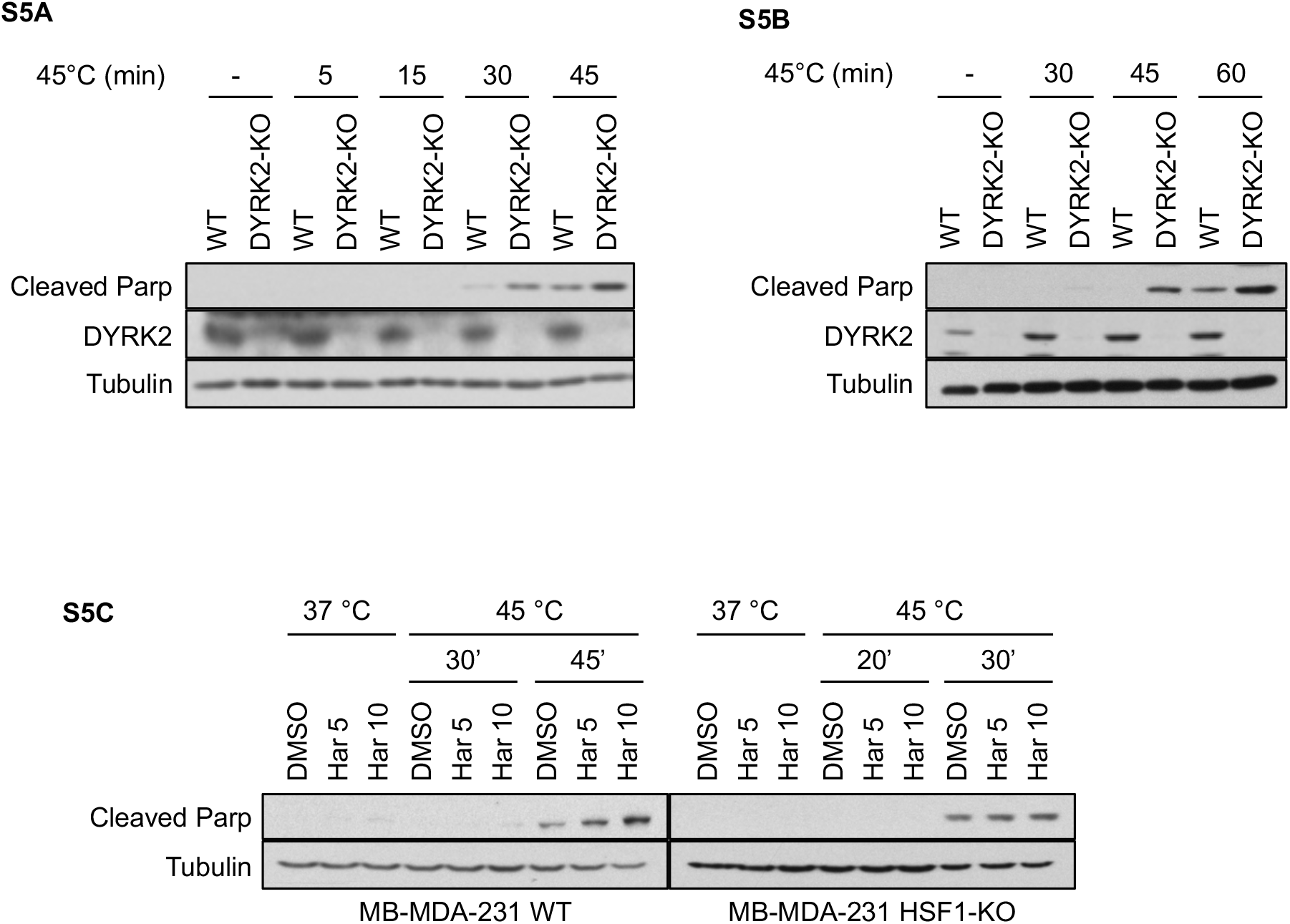
Related to Figure 5. DYRK2 reduces sensitivity to proteotoxic stress via HSF1. **A and B)** Control (WT) and DYRK2-KO MDA-MB-468 cells (A) or 293T cells (B) were incubated at 37 °C or at 45 °C for the indicated times, followed by recovery at 37 °C. On the next day, cells were lysed and the levels of apoptosis were analysed by western blotting using an antibody that recognises cleaved PARP. **C)** Control (WT) *(left panel)* or HSF1-KO *(right panel)* MDA-MB-231 cells were treated with vehicle or increasing concentrations of harmine. After two hours, cells were incubated at 37 °C or at 45 °C for the indicated times, followed by recovery at 37 °C. On the next day, cells were lysed and the levels of apoptosis were analysed by western blotting using an antibody that recognises cleaved PARP.

**Figure S6.**
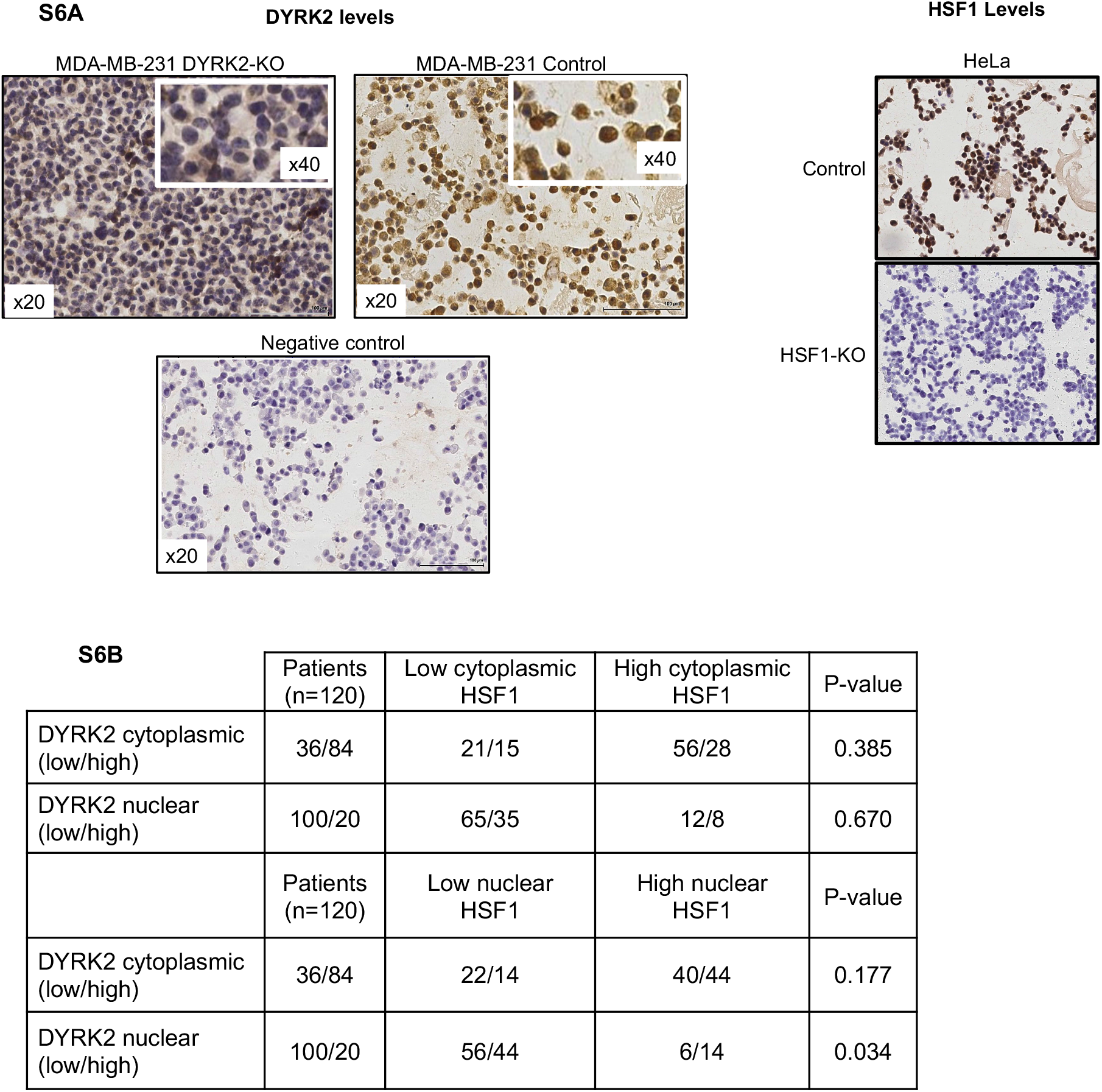
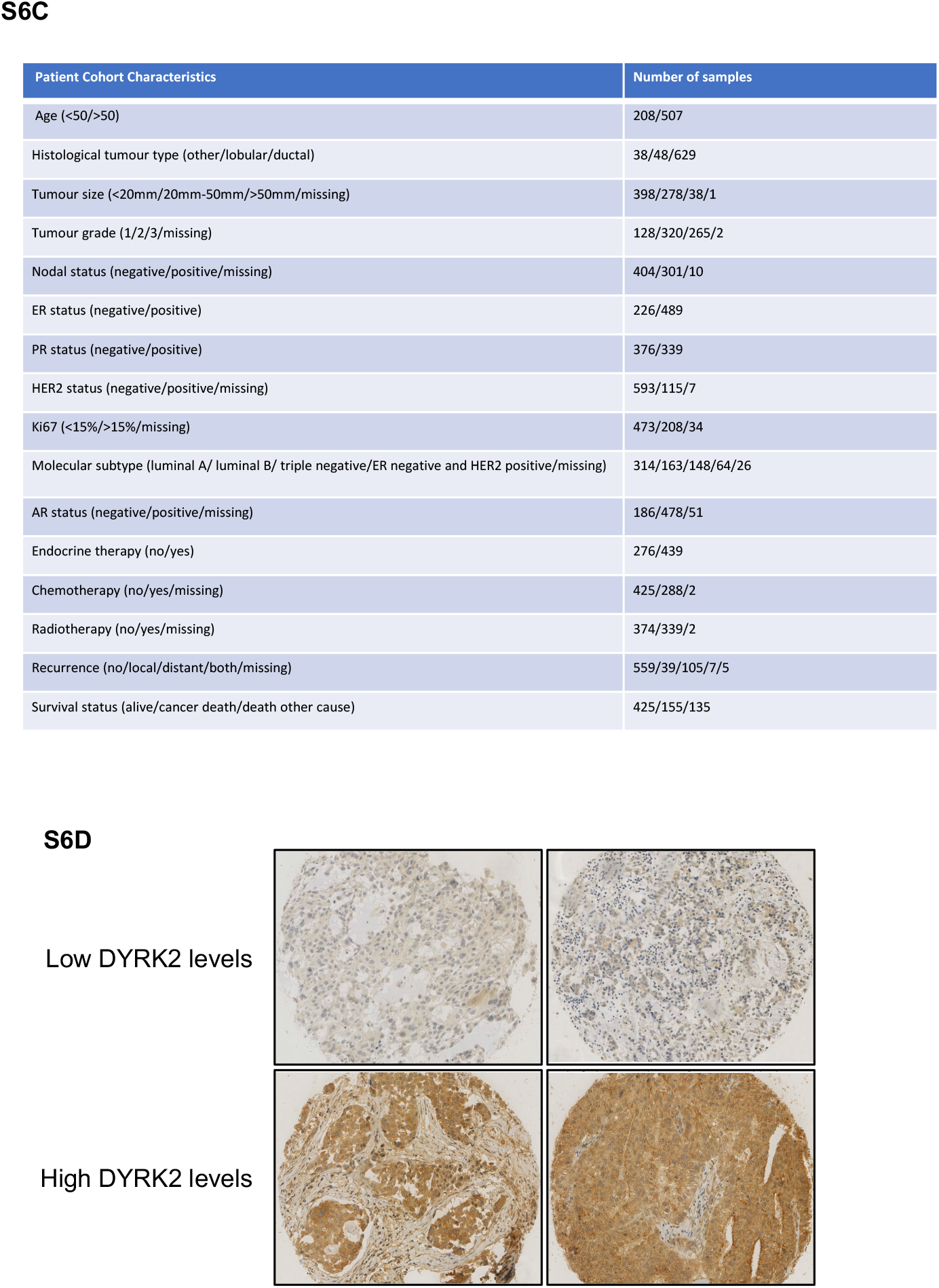
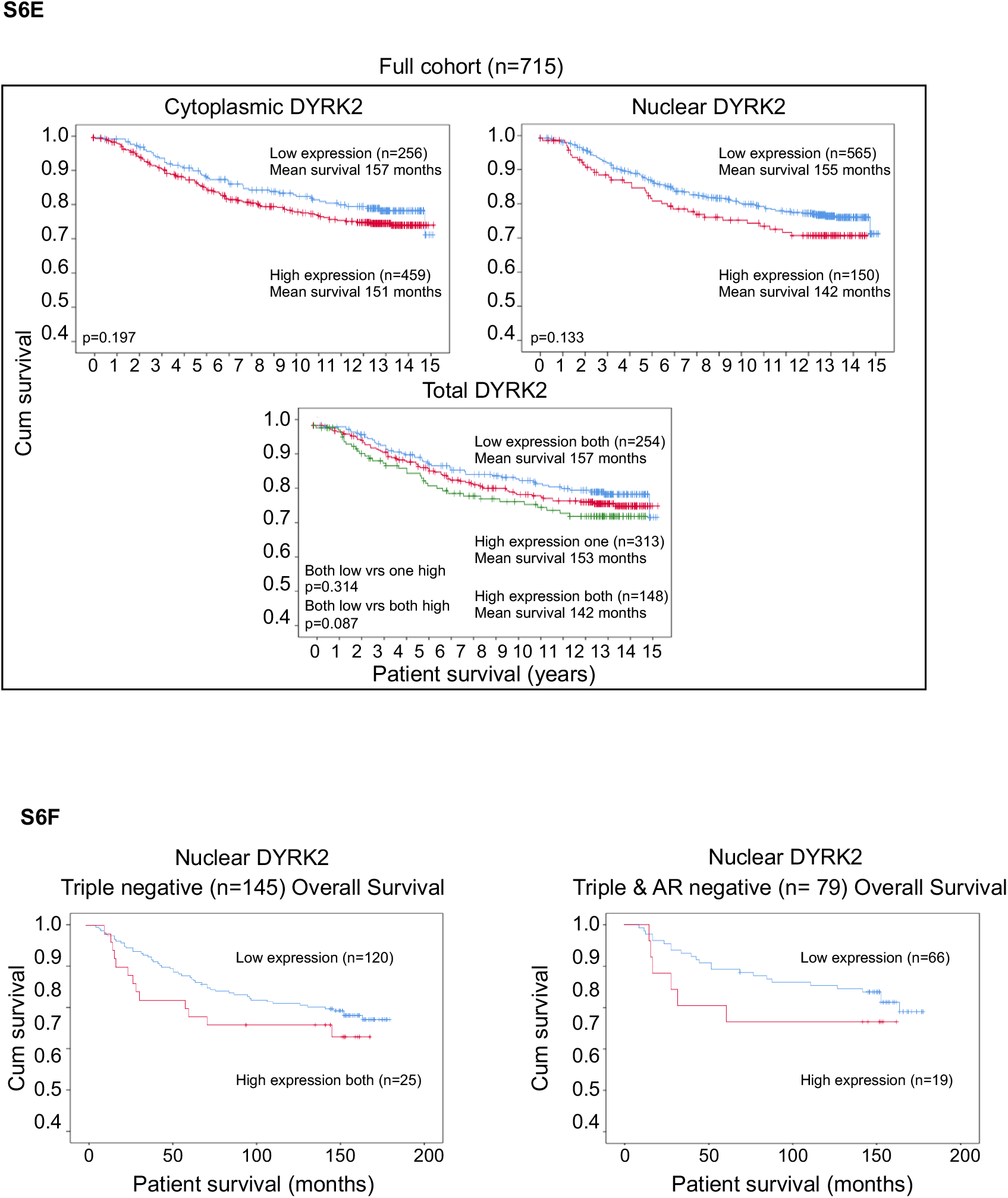
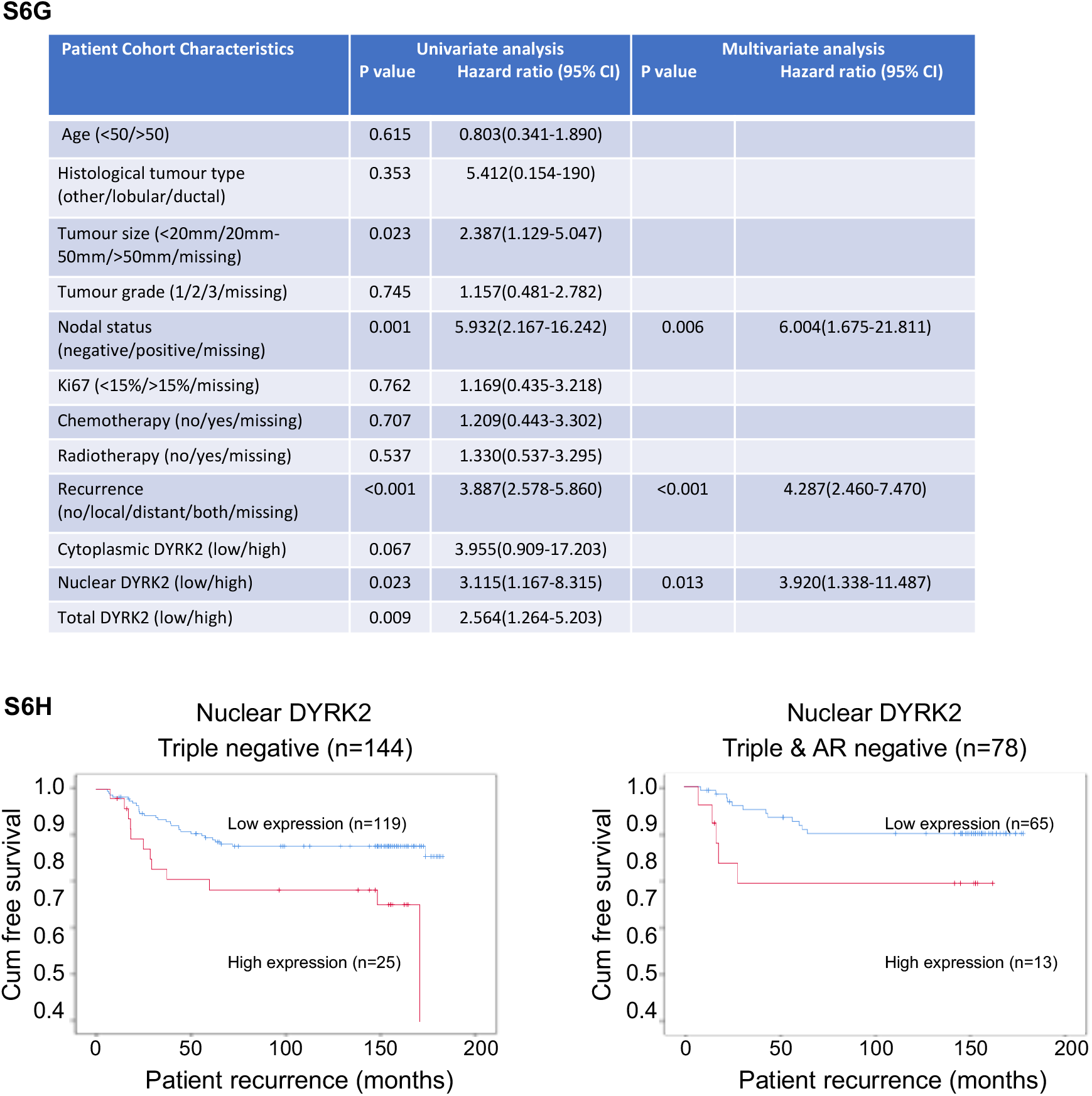
Related to Figure 6. DYRK2 levels correlate with HSF1 nuclear levels, prognosis and tumour recurrence in tissue from TNBC patients. **A)** DYRK2 and HSF1 antibody validation using Formalin-Fixed, Paraffin-Embedded (FFPE) cell pellets. **B)** Chi Square test correlation between low and high HSF1 and DYRK2 protein expression in triple negative breast cancer samples. Significance p< 0.05. **C)** Tissue microarray details. **D)** Representative samples showing high and low DYRK2 staining. **E)** Relationship between cytoplasmic, nuclear and total DYRK2 levels and cancer-specific survival in patients with invasive ductal breast cancer (total cohort). **F)** Relationship between nuclear DYRK2 levels and overall survival in patients with triple negative (left panel) or triple negative and AR negative (right panel) invasive ductal breast cancer. **G)** Univariate and multivariate survival analysis for patients with triple negative and AR negative breast cancer to assess the relationship between clinico-pathological characteristics, DYRK2 and cancer specific survival. **H)** Relationship between nuclear DYRK2 levels in tumour cells and time to recurrence in patients with triple negative (left panel) or triple negative and AR negative (right panel) invasive ductal breast cancer.

